# Whole-brain annotation and multi-connectome cell typing quantifies circuit stereotypy in *Drosophila*

**DOI:** 10.1101/2023.06.27.546055

**Authors:** Philipp Schlegel, Yijie Yin, Alexander S. Bates, Sven Dorkenwald, Katharina Eichler, Paul Brooks, Daniel S. Han, Marina Gkantia, Marcia dos Santos, Eva J. Munnelly, Griffin Badalamente, Laia Serratosa Capdevila, Varun A. Sane, Markus W. Pleijzier, Imaan F.M. Tamimi, Christopher R. Dunne, Irene Salgarella, Alexandre Javier, Siqi Fang, Eric Perlman, Tom Kazimiers, Sridhar R. Jagannathan, Arie Matsliah, Amy R. Sterling, Szi-chieh Yu, Claire E. McKellar, FlyWire Consortium, Marta Costa, H. Sebastian Seung, Mala Murthy, Volker Hartenstein, Davi D. Bock, Gregory S.X.E. Jefferis

## Abstract

The fruit fly *Drosophila melanogaster* combines surprisingly sophisticated behaviour with a highly tractable nervous system. A large part of the fly’s success as a model organism in modern neuroscience stems from the concentration of collaboratively generated molecular genetic and digital resources. As presented in our FlyWire companion paper^1^, this now includes the first full brain connectome of an adult animal. Here we report the systematic and hierarchical annotation of this ∼130,000-neuron connectome including neuronal classes, cell types and developmental units (hemilineages). This enables any researcher to navigate this huge dataset and find systems and neurons of interest, linked to the literature through the Virtual Fly Brain database^2^. Crucially, this resource includes 4,552 cell types. 3,094 are rigorous consensus validations of cell types previously proposed in the “hemibrain” connectome^3^. In addition, we propose 1,458 new cell types, arising mostly from the fact that the FlyWire connectome spans the whole brain, whereas the hemibrain derives from a subvolume. Comparison of FlyWire and the hemibrain showed that cell type counts and strong connections were largely stable, but connection weights were surprisingly variable within and across animals. Further analysis defined simple heuristics for connectome interpretation: connections stronger than 10 unitary synapses or providing >1% of the input to a target cell are highly conserved. Some cell types showed increased variability across connectomes: the most common cell type in the mushroom body, required for learning and memory, is almost twice as numerous in FlyWire as the hemibrain. We find evidence for functional homeostasis through adjustments of the absolute amount of excitatory input while maintaining the excitation-inhibition ratio. Finally, and surprisingly, about one third of the cell types proposed in the hemibrain connectome could not yet be reliably identified in the FlyWire connectome. We therefore suggest that cell types should be defined to be robust to inter-individual variation, namely as groups of cells that are quantitatively more similar to cells in a different brain than to any other cell in the same brain. Joint analysis of the FlyWire and hemibrain connectomes demonstrates the viability and utility of this new definition. Our work defines a consensus cell type atlas for the fly brain and provides both an intellectual framework and open source toolchain for brain-scale comparative connectomics.

## Main

The adult fruit fly represents the current frontier for whole brain connectomics. With 127,978 neurons, the newly completed full adult female brain connectome is intermediate in log scale between the first connectome of *C. elegans* (302 neurons^4, 5^) and the mouse (10^8 neurons), a desirable but currently intractable target^6^. The availability of a complete adult fly brain connectome now allows brain-spanning circuits to be mapped and linked to circuit dynamics and behaviour as has long been possible for the nematode and more recently the *Drosophila* larva (3,016 neurons^7^). However the adult fly has richer behaviour, including complex motor control while walking or in flight^8–10^, courtship behaviour^11, 12^, involved decision making^13, 14^, flexible associative memory^15–21^, spatial learning^22, 23^ and complex^24–28^ multisensory^29–31^ navigation.

The FlyWire connectome reported in our companion paper^1^ is by some margin the largest and most complex yet obtained. The full connectome, derived from the ∼100 teravoxel FAFB whole-brain electron microscopy (EM) volume^32^, can be represented as a graph with 127,978 nodes and ∼16.5M weighted edges. In this paper we formulate and answer a number of key questions essential to interpreting connectomes at this scale: 1) How do we know which edges are important? 2) How can we simplify the connectome graph to aid automated or human analysis? 3) To what extent is this connectome a snapshot of a single brain or representative of this species as a whole; or to put it more provocatively, have we collected a snowflake? These questions are inextricably linked with connectome annotation and cell type identification^33, 34^ within and across datasets.

At the most basic level, navigating this connectome would be extremely challenging without a comprehensive system of annotations, which we now provide. Our annotations provide an indexed and hierarchical human-readable parts list^33, 35^, allowing biologists to explore their systems and neurons of interest. Connectome annotation is also crucial to ensuring data quality since it inevitably reveals segmentation errors that must be corrected. Furthermore, there is a rich history in *Drosophila* of probing the circuit basis of a wide range of innate and learned behaviours as well as their developmental genetic origins; realising the full potential of this dataset is only possible by cross-identifying cell types within the connectome with those characterised in the published and in-progress literature. This paper reports this key component of the connectome together with the open source tools and resources we have generated. Because the annotation and proofreading of the connectome are inextricably linked, Dorkenwald *et al*.^1^ and this paper should be co-cited by users of the FlyWire resource.

Comparison with cell types proposed using the partial hemibrain connectome^3^ confirmed that the majority of fly cell types are highly stereotyped, as well as defining simple rules for which connections within a connectome are reliable and therefore more likely to be functional. However, this comparison also revealed unexpected variability in some cell types and demonstrated that many cell types originally reported in the hemibrain could not be reliably re-identified. This discovery necessitated the development and application of a new robust approach for defining cell types jointly across connectomics datasets. Overall, this effort lays the foundation both for deep interrogation of current and anticipated fly connectomes from normal individuals, but also future studies of sexual dimorphism, experience-dependent plasticity, development and disease at whole brain scale.

## Results

### Hierarchical annotation of a connectome

Annotations defining different kinds of neurons are key to exploring and interpreting any connectome; but with the FlyWire connectome now exceeding the 100,000 neuron mark, they are also both of increased significance and more challenging to generate. We defined a comprehensive, systematic, and hierarchical set of annotations, based on the anatomical organisation of the brain (Figure 1; <u>Supplemental Video 1+2</u>), as well as the developmental origin and coarse morphology of neurons (Figure 2). Building on these as well as validating cell types identified from pre-existing datasets, we then defined a set of consensus terminal cell types intended to capture the finest level of organisation that is reproducible across brains (Figure 3). For the connectome version that we report jointly with Dorkenwald *et al.*^1^ the central brain can be considered a finished connectome by virtue of multiple rounds of synergistic proofreading and annotation; in contrast, the newly proofread optic lobes will likely see further updates through additional rounds of quality control. For this reason, although coarser annotations are provided brainwide, our finest scale cell typing focuses on neurons with arbours in the central brain. Nevertheless this represents most cell types in the fly brain; while intrinsic neurons of the optic lobe are the majority of neurons in the fly brain, their highly parallel architecture means they contain only about 200 cell types^36, 37^, or approximately 20x fewer than the central brain.

**Figure 1:**
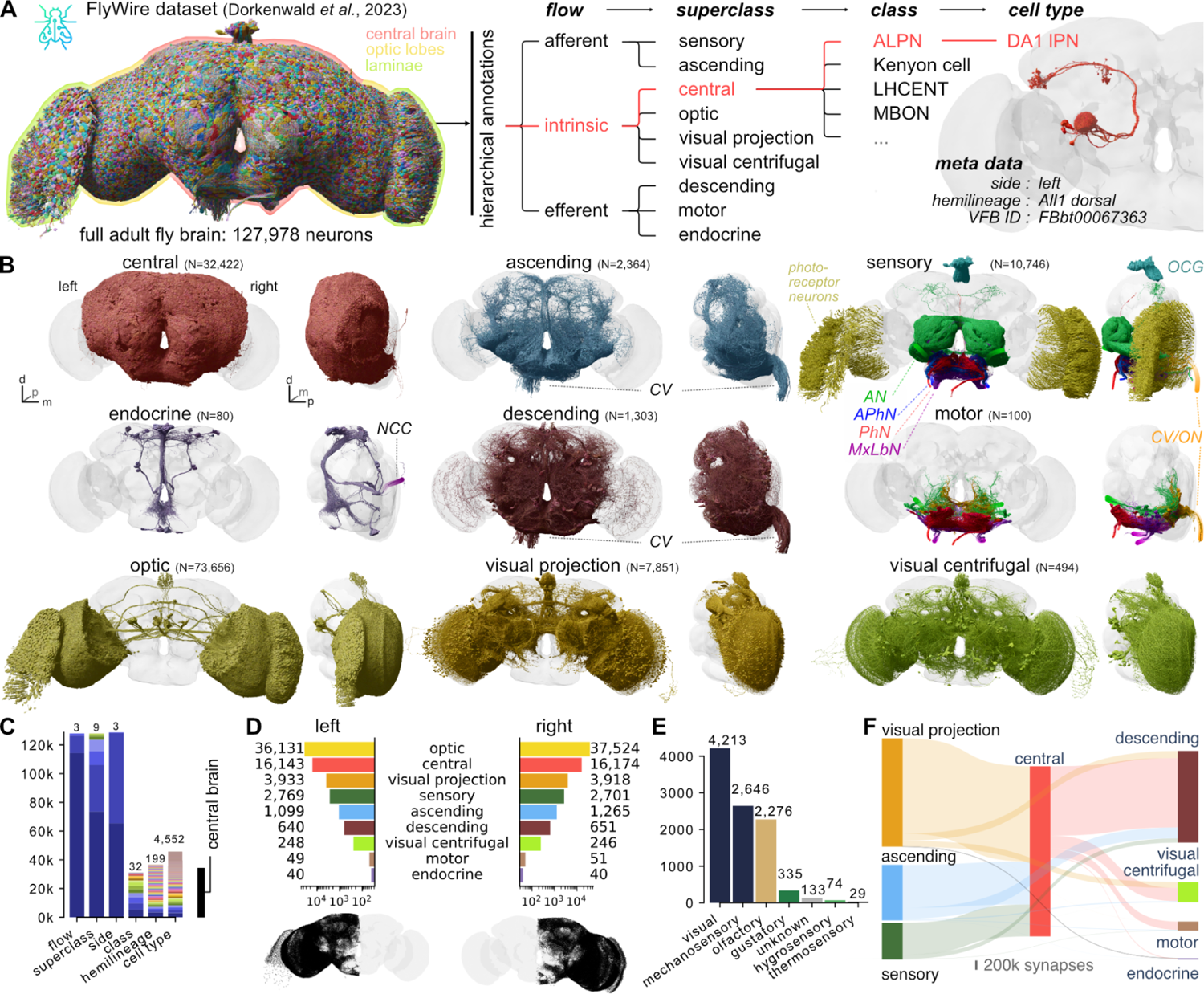
Hierarchical annotation schema for a whole brain connectome. **A** Hierarchical annotation schema for the FlyWire dataset (see companion paper: Dorkenwald *et al*., 2023^1^). **B** Renderings of neurons for each superclass. **C** Annotation counts per field. Each colour within a bar represents discrete values; numbers above bars count the discrete values. **D** Left vs right neuron counts per superclass. “Sensory” count excludes visual (photoreceptor) neurons. Bottom shows left and right soma locations, respectively. **E** Break down of *sensory* neuron counts into modalities. **F** Flow chart of superclass-level, feed-forward (afferent → intrinsic → efferent) connectivity. Abbreviations: CV, cervical connective; AN, antennal nerve; PhN, pharyngeal nerve; APhN, accessory pharyngeal nerve; MxLbN, maxillary-labial nerve; NCC, corpora cardiaca nerves; ON, occipital nerves; OCG, ocellar ganglion.

**Figure 2:**
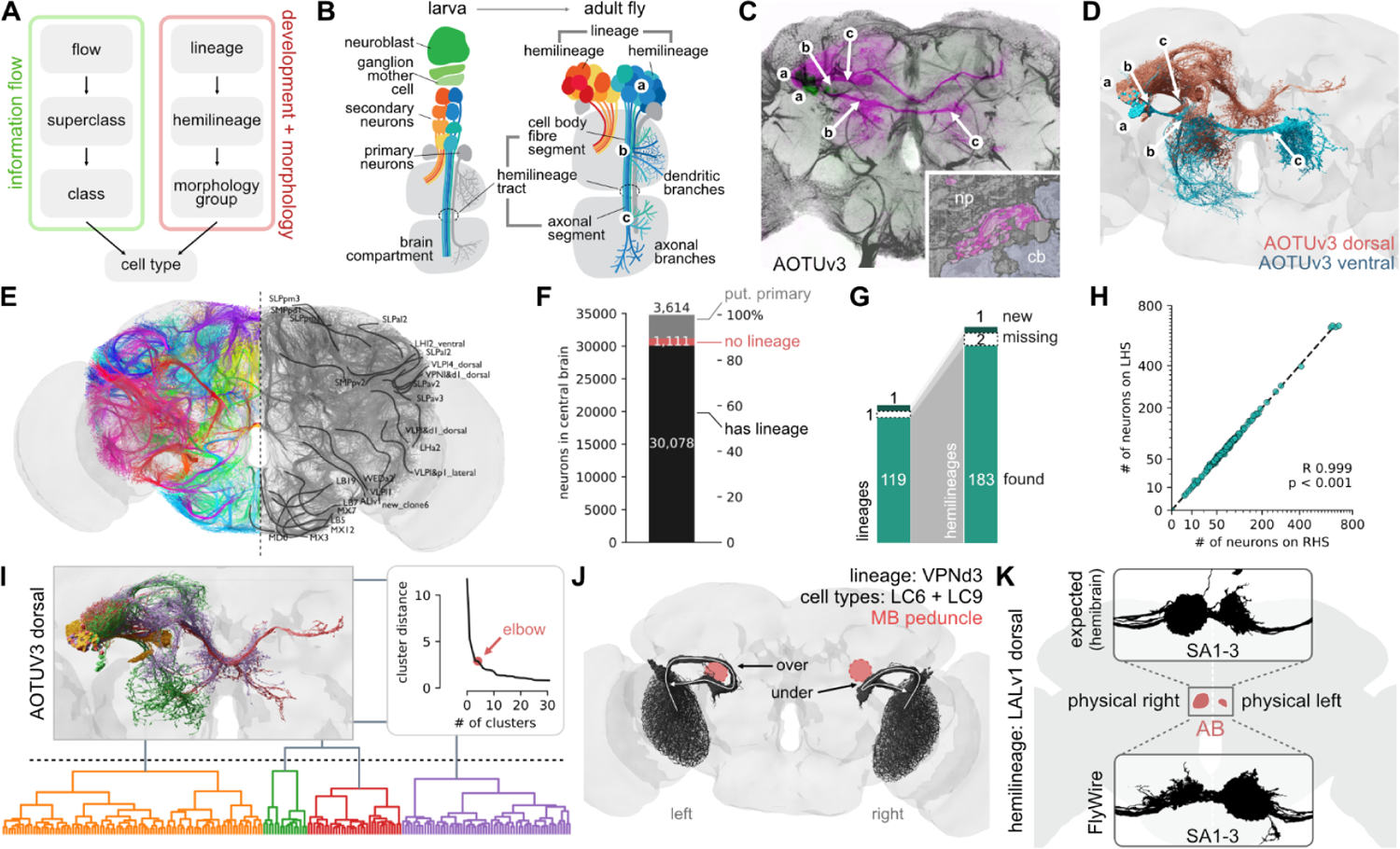
Annotation of developmental units. **A** Illustration of the two complementary sets of annotations. **B** Developmental organisation of neuroblast hemilineages. **C** Light-level image of an example AOTUv3 lineage clone; lower case letters link canonical features of each hemilineage to the cartoon in B. Inset shows cell body fibre tract in the EM. **D** AOTUv3 neurons in Flywire split into its two hemilineages. **E** Cell body fibre bundles from all identified hemilineages, partially annotated on the right. **F** Number of central brain neurons with identified lineage; annotation of (putative) primary neurons are based on literature or expert assessment of morphology. **G** Number of identified unique (hemi-)lineages. **H** Left versus right number of neurons contained in each hemilineage. **I** Morphological clustering of each hemilineage defines subgroups. **J** LC6 and LC9 neurons (lineage VPNd3) of the right and left hemispheres take different routes in FlyWire to equivalent destinations. Mushroom body (MB) peduncle is shown in pink. **K**. The asymmetric body (AB) of the fly is a stereotyped left-right asymmetric structure in the centre of the fly brain. Insets show axons of neuron types SA1, SA2 and SA3, the major input to the AB. FlyWire neurons (bottom) are flipped relative to expectation (top, hemibrain neurons).

**Figure 3:**
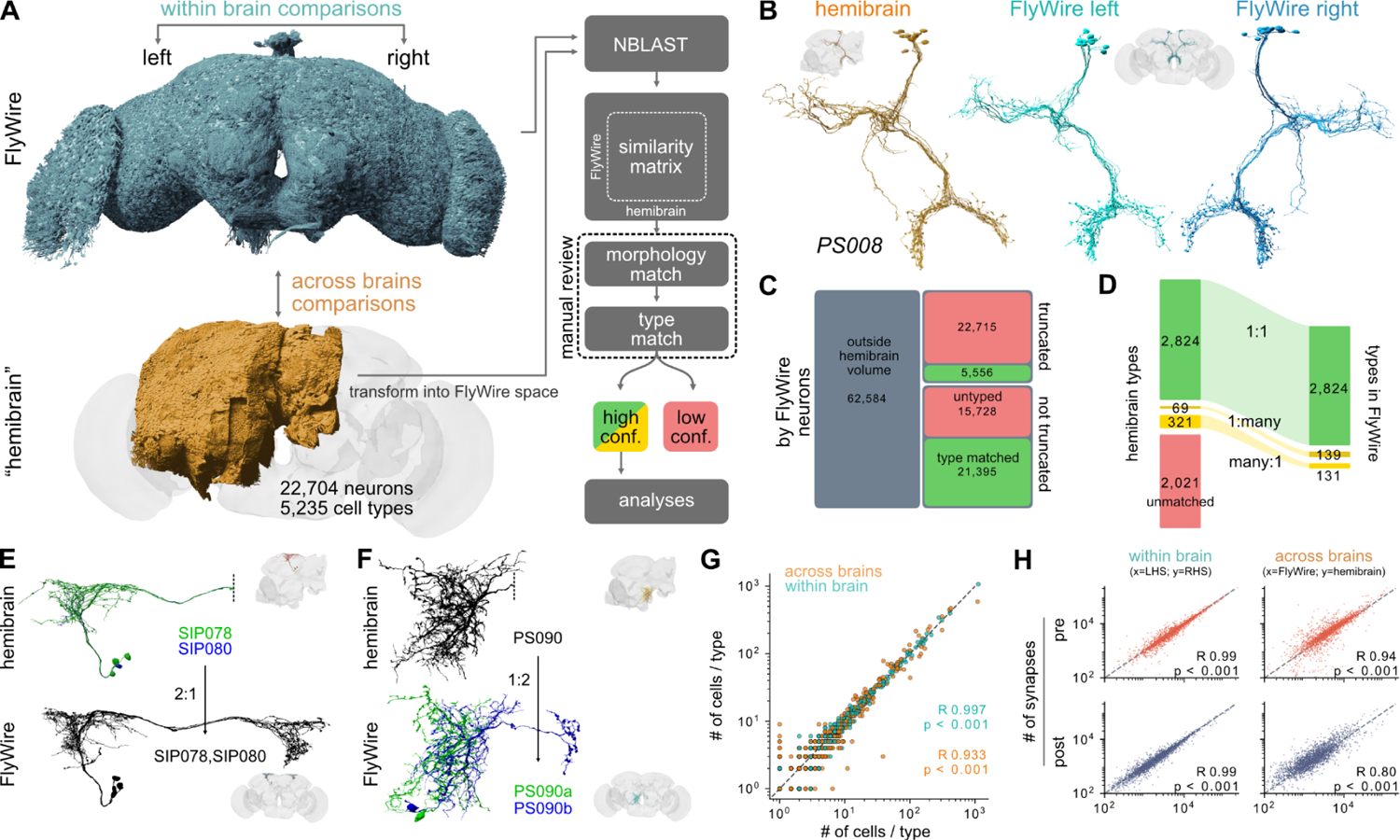
Across-brain stereotypy. **A** Schematic of the pipeline for matching neurons between FlyWire and the Janelia “hemibrain” connectomes. **B** Example for high confidence cell type (PS008) unambiguously identifiable across all three hemispheres. **C** Counts of FlyWire neurons that were assigned a hemibrain type. **D** Number of hemibrain cell types that were successfully identified and the resulting number of FlyWire cell types. **E** Example for a many:1 hemibrain type match. **F** Example for a 1:many cell type match. **G** Number of cells per cross-matched cell type within- (FlyWire left vs right) and across-brains (Flywire vs hemibrain). **H** Number of pre- and post-synapses per cross-matched cell type. *R* is Pearson correlation coefficient.

We first collected and curated basic metadata for every neuron in the dataset including soma position and side, and exit or entry nerve for afferent and efferent neurons (Figure 1). Our group also predicted neurotransmitter identity for all neurons as reported elsewhere^38^. We then defined a hierarchy of four levels: *flow* → *superclass* → *class* → *cell type* which provide salient labels at different granularities (Figure 1A; Supplemental Table S1; <u>Figure S1</u>).

The first two levels, *flow* and *superclass*, were densely annotated: every neuron is either afferent, efferent or intrinsic to the brain (*flow*) and falls into one of the 9 *superclasses*: sensory (periphery→brain), motor (brain→periphery), endocrine (brain→corpora allata/cardiaca), ascending (ventral nerve cord→brain), descending (brain→ventral nerve cord), visual projection (optic lobes→central brain), visual centrifugal (central brain→optic lobes), or intrinsic to the optic lobes or the central brain (Figure 1B, Supplemental Table S2), A mapping to the virtualflybrain.org^2^ database allows cross-referencing neuron/types with other publications (see Methods for details).

The *class* field contains pre-existing neurobiological groupings from the literature (e.g. for central complex neurons; see Supplemental Table S3) and is sparsely annotated (40% of the central brain), in large part because past research has favoured some brain areas over others. Finally, over half (57%) of the neurons associated with the central brain (i.e. both inputs to the central brain as well as intrinsic neurons with their cell bodies in the central brain) were given a terminal *cell type* most of which could be linked to at least one report in the literature (Figure 1C). The remaining neurons are typeable even if they cannot be unambiguously linked to previously reported cell types, and we provide a path for this below (“<u>Toward multi-connectome cell typing</u>”). In total, we collected over 730k annotations for all 127,978 neurons; all are available for download and 580k are already visible in codex.flywire.ai under the Classification heading for each cell. 32,422 (28%) neurons are intrinsic to the central brain and 73,656 (56%) neurons are intrinsic to the optic lobes. The optic lobes and the central brain are connected via 7,851 visual projection and 494 visual centrifugal neurons. The central brain receives afferent input via 5,495 sensory and 2,364 ascending neurons. Efferent output is realised via 1,303 descending, 80 endocrine and 100 motor neurons.

We find striking stereotypy in the number of central brain intrinsic neurons: e.g. between the left and the right hemisphere they differ by only 42 (0.1%) neurons. For superclasses with less consistency in left vs right counts such as the ascending neurons (166, 7%) the discrepancies are typically due to ambiguity in the sidedness (Figure 1D, see <u>Methods</u> for details).

Combining the dense superclass annotation for all neurons with the connectome^1^ gives a birds-eye view of the input/output connectivity of the central brain (Figure 1F): 56% percent of the central brain’s synaptic input comes from the optic system; 25% from the ventral nerve cord (VNC) via ascending neurons; and only 19% from peripheral sensory neurons. This is somewhat surprising since sensory neurons are almost as numerous as visual projection neurons (Figure 1D,E); individual visual projection neurons therefore provide about 2.5 times more synapses underscoring the value of this information stream. Input neurons make about two synapses onto central brain neurons for every one synapse onto output neurons. Most output synapses target the VNC via descending neurons (77%); the rest provide centrifugal feedback onto the optic system (13%), motor neuron output (9%), and endocrine output to the periphery (1%).

### A full atlas of neuronal lineages in the central brain

Our top-level annotations (flow, superclass, class) provide a systematic but relatively coarse grouping of neurons compared with > 5,000 terminal cell types expected from previous work on the hemibrain^3^. We therefore developed an intermediate level of annotation based on *hemilineages* – this provides a powerful bridge between the developmental origin and molecular specification of neurons and their place within circuits in the connectome (Figure 2A).

Central brain neurons and a minority of visual projection neurons are generated by ∼120 identified neuroblasts per hemisphere. Each of these stem cells is defined by a unique transcriptional code and generates a stereotyped lineage in a precise birth order by asymmetric division (Figure 2B)^39–42^. Each neuroblast typically produces two hemilineages^43, 44^ that differ markedly in neuronal morphology and can express different neurotransmitters from one another, but neurons in each hemilineage usually express a single fast-acting transmitter^38, 45^. Hemilineages therefore represent a natural functional as well as developmental grouping by which to study the nervous system. Within a hemilineage, neurons form processes that extend together in one cohesive bundle (the hemilineage tract) which enters, traverses, and interconnects neuropil compartments in a stereotypical pattern (Figure 2C). Comparing these features between EM and prior light-level data^46–49^ allowed us to compile the first definitive atlas of all hemilineages in the central brain (Figure 2C-E; see <u>Methods</u> for details).

In total, we successfully identified 120 neuroblast lineages in FlyWire comprising 183 hemilineages for 30,078 (86.4%) central brain neurons (Figure 2F; <u>Figure S2.1</u>). The majority of the unassigned neurons are likely primary neurons born during embryonic development, which account for 10% of neurons in the adult brain^50, 51^. We tentatively designated 3,614 (10.4%) as primary neurons either based on specific identification in the literature^42^ or expert assessment of diagnostic morphological features such as broader projections and larger cell bodies. A further 1,111 (3.2%) neurons did not co-fasciculate with any hemilineage tracts even though their morphology suggested that they are later-born secondary neurons^52^. This developmental atlas is comprehensive since all but one (SLPpm4^48^) of the 120 previously described lineages were identified, in addition to one new lineage (LHp3/CP5). We were able to find the expected hemilineages in all but two cases (LHa1_posterior and SLPal4_anterior; Figure 2G). The number of neurons per hemilineage can vary widely (Figure 2H): for example, counting both hemispheres FLAa1 contains just 30 neurons while MBp4 (which makes the numerous Kenyon cells required for memory storage) has 1,335. In general, however, the number of neurons per hemilineage is between 31 and 113 (10/90 percentile, respectively). Nevertheless, the numbers of neurons within each hemilineage was highly reliable differing only by 3% (+/-4) between left and right hemispheres (Figure 2G, right). This is consistent with the near-equality of neurons per hemisphere noted in Figure 1, and indicates great precision in the developmental programs controlling neuron number.

Although hemilineages typically contain functionally and morphologically related neurons, subgroups can be observed^53^. We further divided each hemilineage into “morphology groups” each innervating similar brain regions and taking similar internal tracts using NBLAST morphological clustering^54^ (Figure 2I, <u>Figure S2.1</u>, <u>Supplemental Files 2 and</u> <u>3</u>; <u>Supplemental Video 3</u>). This generated a total of 681 groupings which provide an additional layer of annotations between the hemilineage and cell type levels.

The fly brain is mostly left-right symmetric but inspection of the FlyWire dataset reveals some large asymmetries. For example the visual projection neurons of the VPNd3 lineage^48^ (including cell types LC6 and LC9) form a large axon bundle. This bundle follows the normal path in the right hemisphere^55^ but in the left hemisphere it loops over (i.e. medial) the mushroom body peduncle; nevertheless, the axons still find their correct targets (Figure 2J). Besides such unexpected asymmetries, the fly brain contains one structure, the asymmetric body (AB), that is reproducibly ∼4 times larger on the right hemisphere of the fly’s brain than on the left^56–58^. Surprisingly, we found that the left/right volume of the AB appeared inverted in FlyWire (Figure 2K). The cause of this is not biological but rather because the whole brain volume was accidentally flipped during acquisition of the original FAFB EM image data which underlies FlyWire (<u>Figure S2.2A</u> and Methods section <u>FAFB</u> <u>Laterality</u>). Note that our “*side*” annotation and all references to left or right in this paper refer to the correct side of the fly’s brain.

### Validating cell types across brains

Next, we sought to compare FlyWire against the hemibrain connectome^3^; this contains most of one central brain hemisphere and parts of the optic lobe. The hemibrain was previously densely cell-typed by a combination of two automated procedures followed by extensive manual review^3, 59–61:^ NBLAST morphology clustering initially yielded 5,235 morphology types; multiple rounds of CBLAST connectivity clustering split some types, generating 640 connectivity types for a final total of 5,620 types. We have reidentified just 3% of connectivity types and therefore use the 5,235 morphology types as a baseline for comparison. Although 512 (10%) of the hemibrain cell types were previously established in the literature and recorded in virtualflybrain.org^2^, principally through analysis of genetic driver lines^34^, the great majority (90%) were newly proposed using the hemibrain, i.e. derived from a single hemisphere of a single animal. This was reasonable given the pioneering nature of the hemibrain reconstruction, but the availability of the FlyWire connectome now allows for a more stringent reexamination.

We approach this by considering each cell type in the hemibrain as a prediction: if we can reidentify a distinct group of cells with the same properties in both hemispheres of the FlyWire dataset, then we conclude that a proposed hemibrain cell type has been tested and validated. To perform this validation, we first used non-rigid 3D registration to map 3D meshes and skeletons of all hemibrain neurons into FlyWire space, enabling direct 3D co-visualisation of both datasets and a range of automated analyses (Figure 3A-B). We then used NBLAST^54^ to calculate morphological similarity scores between all hemibrain neurons and the ∼62.4k FlyWire neurons with extensive arbours within the hemibrain volume (<u>Figure</u> <u>S3A-C</u>). Candidate matches were manually reviewed by co-visualisation and only those with high confidence were accepted (Figure 3C-D, see <u>Methods</u> for additional details).

The majority of hemibrain cell types (61%; 3,214 of 5,235 types, Figure 3D) were found in the FlyWire dataset. We started with exclusively morphological matching, prioritising neurons for which the top NBLAST scores were clearly distinct from the next most similar neurons. Crucially, the initial morphological matching process generated a large corpus of shared labels between datasets; with these in place we developed a novel across-dataset connectivity clustering method which allowed us to investigate and resolve difficult cases (see Methods – <u>hemibrain cell type matching with connectivity</u>). 4% of proposed hemibrain types were combined to define new, “composite” types (e.g. “SIP078,SIP080”) because the hemibrain split could not be recapitulated when examining neurons from both FlyWire and the hemibrain (Figure 3E; <u>Figure S3D,E</u>). This is not too surprising as the hemibrain philosophy was explicitly to err on the side of splitting in cases of uncertainty^3^. We found that 1% of proposed hemibrain types needed to be split, for example because truncation of neurons in the hemibrain removed a key defining feature (Figure 3F). Together these revisions mean that the 3,214 reidentified hemibrain cell types map onto 3,094 consensus cell types (Figure 3D). All revisions were confirmed by across-dataset connectivity clustering. Surprisingly, 2,021 (39%) of hemibrain cell types could not yet be re-identified in FlyWire. Ambiguities due to hemibrain truncation can partially explain this: we were much more successful at matching neurons that were not truncated in the hemibrain (Figure 3C). However this appears to not to be the main explanation. Further investigation (Figure 6) suggests that although, with time, we and our colleagues should be able to validate a minority of these unmatched hemibrain types, the majority are not exactly replicable across animals. Instead we show that multi-connectome analysis can generate cell type proposals that are robust to inter-individual variation.

In conclusion, we validated 3,094 high-confidence consensus cell type labels for 43,600 neurons from three different hemispheres and two different brains (e.g. Figure 3B; <u>Figure S8</u>). Collectively these cross-matched neurons cover 39.7% of central brain edges (comprising 43% of synapses) in the FlyWire graph. This body of high-confidence cross-identified neurons enables both within- (FlyWire left versus right hemisphere) and across-brain (FlyWire vs hemibrain) comparisons.

### Cell types are highly stereotyped within and across brains

Using the consensus cell type labels, we find that the numbers of cells per type across the three hemispheres are closely correlated (Figure 3G). About 1 in 6 cell types shows a difference in numbers between the left and right hemisphere and 1 in 4 across brains (FlyWire versus hemibrain). The mean difference in number of cells per type is small though: 0.3 (+/-1.7) within- and 0.7 (+/-10) across brains. Importantly, cell types with fewer neurons per type are less variable (<u>Figure S3H</u>). At the extreme, “singleton” cell types account for 65% of all types in our sample; they often appear to be embryonic-born, or early secondary neurons, and only very rarely comprise more than one neuron: only 2% of neurons that are singletons in both FlyWire hemispheres have more than 1 member in the hemibrain. In contrast, more numerous cell types are also more likely to vary in number both within but even more so across brains (<u>Figure S3H,I</u>).

Synapse counts were also largely consistent within cell types, both within and across brains (Figure 3H). To enable a fair comparison, the FlyWire synapse cloud was restricted to the smaller hemibrain volume. Although this does not correct for other potential confounds such as differences in the synaptic completion rates or synapse detection, pre- and postsynapse counts per cell type were highly correlated, both within- (Pearson R 0.99, p<0.001) and across-brains (Pearson R 0.95/0.8 for pre- and postsynapses, respectively, p<0.001; Figure 3H; <u>Figure S3J,K</u>). This is an important quality control and pre-requisite for subsequent connectivity comparisons.

### Interpreting connectomes

Brain wiring develops through a complex and probabilistic developmental process^62, 63^. To interpret the connectome it is vital to obtain a basic understanding of how variable that biological process is. This is complicated by the fact that the connectome we observe is shaped not just by biological variability but also by technical noise, e.g. from segmentation issues, synapse detection errors and synaptic completion rates (the fraction of synapses attached to proofread neurons) (Figure 4A). Here, we use the consensus cell types to assess which connections are reliably observed across three hemispheres of connectome data. We use the term “edge” to describe the set of connections between two cell types, and its “weight” as the number of unitary synapses forming that connection.

**Figure 4:**
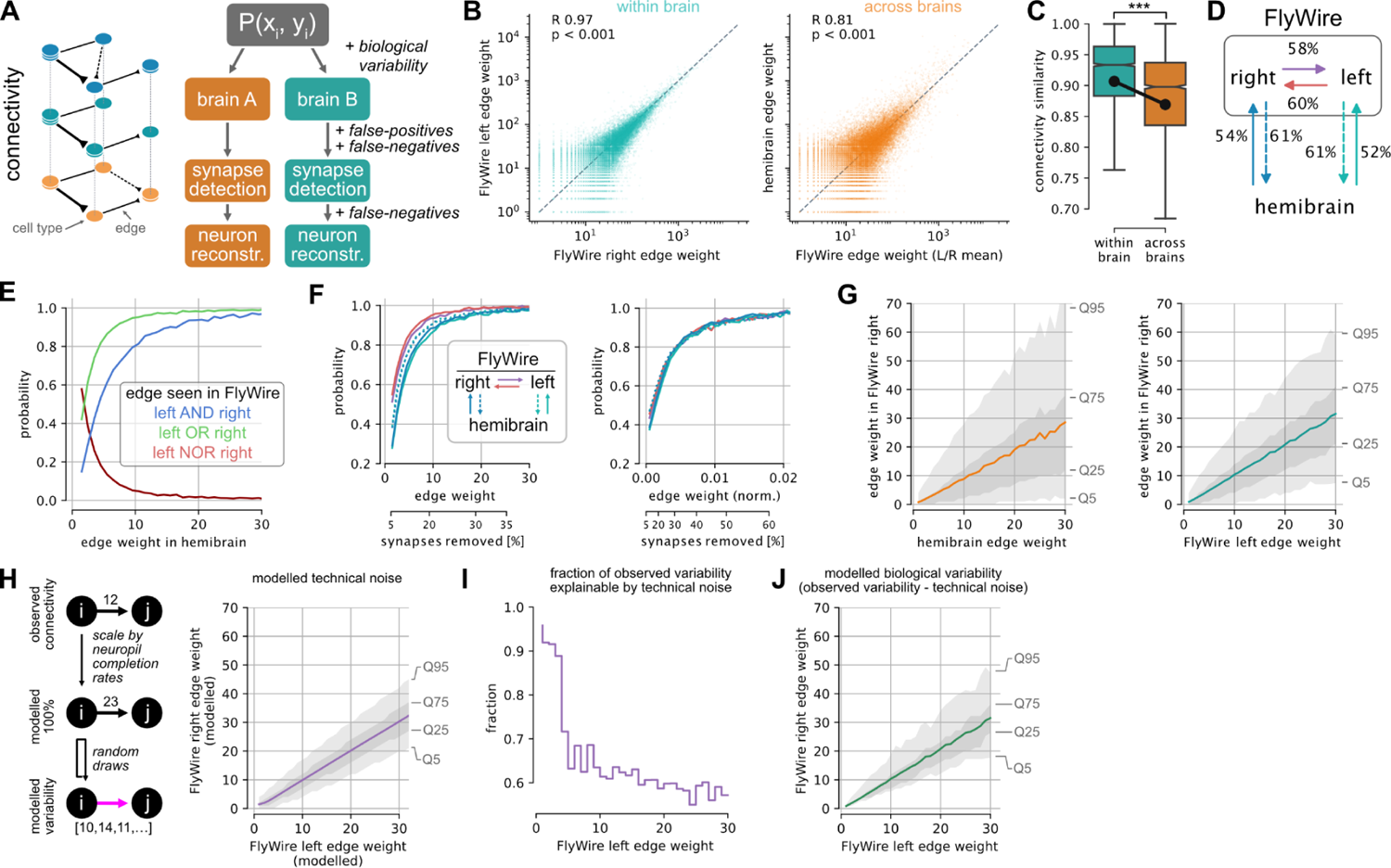
Connectivity stereotypy. **A** Connectivity comparisons and potential sources of variability. **B,C** Edge weights (B) and cosine connectivity similarity (C) between cross-matched cell types. R is Pearson correlation coefficient; ***, p<0.001. **D** Percentage of edges in one hemisphere that can be found in another hemisphere. **E** Probability that an edge present in the hemibrain is found in one, both or neither of the hemispheres in FlyWire. See <u>Figure S4D</u> for plot with normalised edge weights. **F** Probability that edge is found within and across brains as a function of total (left) and normalised (right) edge weight. Second x-axis shows percent of synapses below a given weight. **G** Correlation of across (left) and within (right) edge weights. Envelopes represent quantiles. **H** Model for the impact of technical noise (synaptic completion rate, synapse detection) on synaptic weight from cell types *i* to *j*. The raw weight from the connectome for each individual edge is scaled up by the computed completion rate for all neurons within the relevant neuropil; random draws of the same fraction of those edges then allow an estimate of technical noise. **I** Fraction of FlyWire left/right edge pairs that fall within the 5-95% quantiles for the modelled technical noise. **J** Modelled biological variability.

Weights of individual edges are highly correlated within (Pearson R 0.96, p<0.001) and across (Pearson R 0.81, p<0.001) brains (Figure 4B, <u>Figure S4A</u>). Consistent with this, cell types exhibit highly similar connectivity within as well as across brains (Figure 4C, <u>Figure</u> <u>S4B-C</u>). While the connectivity (cosine) similarity across brains is lower than within brains (p<0.001), the effect size is small (0.033 +/- 0.119) and is at least in part due to the aforementioned truncation in the hemibrain.

Given an edge between two cell types in one hemisphere, what are the odds of finding the same connection in another hemisphere or brain? Examination of 434,790 edges present in at least one of the three brain hemispheres showed that 53% of the edges observed in the hemibrain were also found in FlyWire. This fraction is slightly higher when comparing between the two FlyWire hemispheres: left→right: 57%; right→left: 59% (Figure 4D). Weaker edges were less likely to be consistent: an edge consisting of a single synapse in the hemibrain has a 41% chance to be also present in a single FlyWire hemisphere, and only a 14% chance to be seen in both hemispheres of FlyWire (Figure 4E). In contrast, edges of >10 synapses can be reproducibly found (>90%). Although only 16% of all edges meet this threshold, they comprise ∼79% of all synapses (Figure 4F; <u>Figure S4E</u>). We also analysed normalised edge weights expressed as a fraction of the input onto each downstream neuron; this accounts for the small difference in synaptic completion rate between FlyWire and the hemibrain. With this treatment, the distributions are almost identical for within and across brain comparisons (Figure 4F, compare left and right panels); edges constituting >1.1% of the target cell type’s total inputs have a greater than 90% chance of persisting (Figure 4F, right panel). Around 7% of edges, collectively containing over half (54%) of all synapses, meet this threshold.

We observed that the fraction of edges persisting across datasets plateaued as the edge weight increased. Using a level of 99% edge persistence, we can define a second principled heuristic: edges greater than 40 synapses or 3% edge weight can be considered strong. It is important to note that these statistics defined across the whole connectome can have exceptions in individual neurons. For example, descending neuron DNp42 receives 34 synapses from PLP146 in FlyWire right, but none on the left or hemibrain; this may well be an example of developmental noise (i.e., bona fide biological variability, rather than technical noise).

So far we have only asked a binary question: does an edge exist or not? However, the conservation of edge weight is also highly relevant for interpreting connectomes. Given that an edge is present in two or more hemispheres, what are the odds that it will have a similar weight? Although edge weights within and across brains are highly correlated (Figure 4B), a 30-synapse edge in the hemibrain, for example, will on average consist of 26 synapses in FlyWire, likely because of differences in synaptic detection and completion rates for these two datasets imaged with different EM modalities^1^. The variance of edge weights is considerable: 25% of all 30-synapse hemibrain edges will consist of fewer than 10 synapses in FlyWire, and 5% will consist of only 1-3 synapses. Consistency is greater when looking within FlyWire: a 30-synapse edge on the left will, on average, also consist of 30 synapses on the right. Still, 25% of all 30-synapse edges on the left will consist of 15 synapses or less on the right, and 5% of only 1-7 synapses (Figure 4G).

How much of this edge weight variability is biological and how much is technical? To assess this, the impact of technical noise on a fictive ground truth model was assessed (Figure 4H; <u>Methods</u>). This model was randomly subsampled according to postsynaptic completion rate (in the mushroom body, for example, this ranges from 46% in FlyWire right to 77% in hemibrain; see <u>Figure S4F</u>), and synapses were randomly added and deleted according to the false-positive and -negative rates reported for the synapse detection^64^. Repeated application of this procedure generated a distribution of edge weights between each cell type pair expected due to technical noise alone. On average, 65% of the observed variability of edge weight between hemispheres fell within the range expected due to technical noise; this fraction approached 100% for weaker synapses (Figure 4I). For example, cell type LHCENT3 targets LHAV3g2 with 30 synapses on the left but only 23 on the right of FlyWire, which is within the 5-95% quantiles expected due to technical noise alone. Overall, this analysis shows that observed variability (Figure 4G, right) is greater than can be accounted for by technical noise, establishing a lower bound for likely biological variability (Figure 4J), and suggests another simple heuristic: differences in edge weights of +/-30% or less may be entirely due to technical noise and should not be over-interpreted.

### Variability in the mushroom body

The comprehensive annotation of cell types in the FlyWire dataset revealed that the number of Kenyon cells (KCs), the intrinsic neurons of the MB, is 30% larger per hemisphere than in the hemibrain (2,597 KCs in FlyWire right; 2,580 in FlyWire left; and 1,917 in hemibrain), well above the average variation in cell counts (5% +/-12%). While these KC counts are within the previously reported range^65^, the difference presents an opportunity to investigate how connectomes accommodate perturbations in cell count. The mushroom body contains 5 principal cell classes: KCs, mushroom body output neurons (MBONs), modulatory neurons (DANs, OANs), the DPM and APL giant interneurons^66^ (Figure 5A). KCs further divide into 5 main cell types based on which parts of the mushroom body they innervate: KCab, KCab-p, KCg-m, KCa’b’ and KCg-d (Figure 5B). Of those, KCab, KCa’b’ and KCg-m are the primary recipients of largely random^59, 67^ (but see ^68^) olfactory input via ∼130 antennal lobe projection neurons (ALPNs) comprising 58 canonical types^59, 60^. Global activity in the mushroom body is regulated via an inhibitory feedback loop mediated by APL, a single large GABAergic neuron^69^. Analogous to the mammalian cerebellum, KCs transform the dense overlapping odour responses of the early olfactory system into sparse non-overlapping representations which enable the animal to discriminate individual odours during associative learning^70, 71^. The difference in cell counts is not evenly distributed across all KC types: KCg-m (and to a lesser extent KCg-d and KCa’b’) are almost twice as numerous in FlyWire versus hemibrain while KCab and KCab-p are present in similar numbers (Figure 5C). Protein starvation during the larval stage can induce specific increases in KCg-m number^72^, suggesting that environmental variations in food resources may have contributed to this difference.

**Figure 5:**
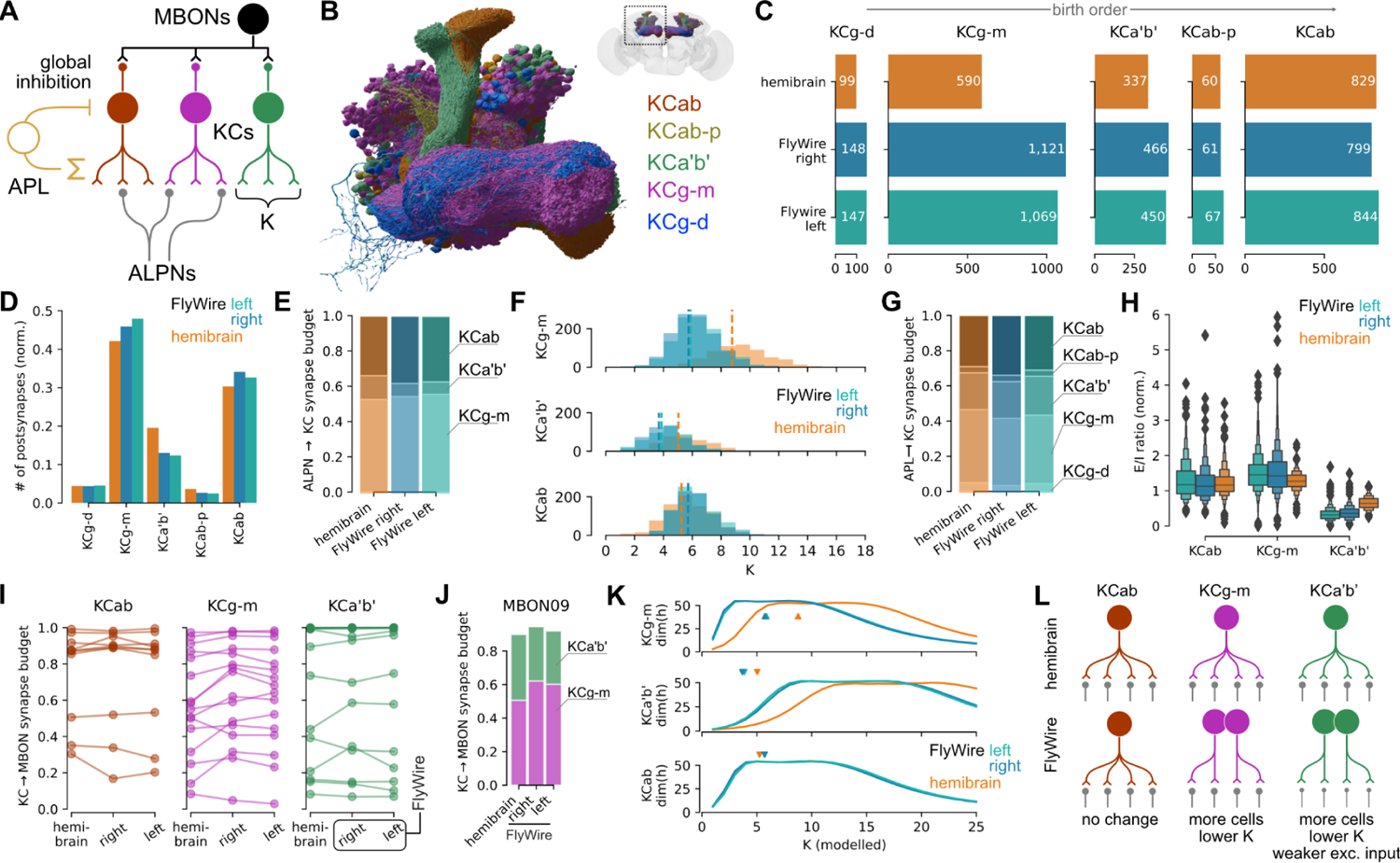
Variability in the mushroom body. **A** Schematic of mushroom body circuits. *K* refers to the number of ALPN types a KC samples from. Neuron types not shown: DANs, DPM, OANs. **B** Rendering of KC cell types. **C** Per-type KC counts across the three hemispheres. **D** KC postsynapse counts, normalised to total KC postsynapses in each dataset. **E** Fractions of ALPN→KC budget spent on individual KC types. **F** Number of ALPN types a KC receives input from, *K*. Dotted vertical lines represent mean. **G** Fraction of APL→KC budget spent on individual KC types. **H** Normalised excitation/inhibition ratio for KCs. **I** Fraction of MBON input budget coming from KCs. Each line represents an MBON type. **J** MBON09 as an example for KC→MBON connectivity. See <u>Figure S5</u> for all MBONs. **K** Dimensionality as function of a modelled *K*. Arrowheads mark observed mean *K*’s. **L** Summarising schematic. Abbreviations: Kenyon cell, KC; ALPN, antennal lobe projection neuron; MBON, mushroom body output neuron; DAN, dopaminergic neuron; APL, anterior paired lateral neuron; OAN, octopaminergic neurons.

How does this affect the mushroom body circuitry? We opted to compare the fraction of the input or output synaptic budget across different KCs since this is well matched to our question and naturally handles a range of technical noise issues that seemed particularly prominent in the mushroom body completion rate (see <u>Methods</u>; <u>Figure S5A</u>). We found that despite the large difference in KCg-m cell counts between FlyWire and hemibrain, this cell type consistently makes and receives 32% and 45% of all KC pre- and postsynapses, respectively (Figure 5D and <u>S5E</u>). This suggested that individual FlyWire KCg-m neurons receive fewer inputs and make fewer outputs than their hemibrain counterparts. The share of ALPN outputs allocated to KCg-m is ∼55% across all hemispheres (Figure 5E), and the average ALPN→KCg-m connection is comparable in strength across hemispheres (<u>Figure</u> <u>S5F</u>); however, each KCg-m receives input from a much smaller number of ALPN types in FlyWire than in the hemibrain (5.74, 5.88 and 8.76 for FlyWire left, right and hemibrain, respectively; Figure 5F). FlyWire KCg-m therefore receive inputs with the same strength but from fewer ALPNs.

This pattern holds for other KCg-m synaptic partners as well. Similar to the excitatory ALPNs, the share of APL outputs allocated to KCg-m is essentially constant across hemispheres (Figure 5G). Therefore each individual KCg-m receives proportionally less inhibition from the APL, as well as less excitation, maintaining a similar excitation-inhibition ratio (Figure 5H). Further, as a population, KCg-m contributes similar amounts of input to MBONs (Figure 5I,J and <u>S3H</u>).

Past theoretical work has shown that the number (*K*) of discrete odour channels (i.e., ALPN types) providing input to each KC has an optimal value for maximising dimensionality of KC activity, and therefore, discriminability of olfactory input^70, 71^. The smaller value for *K* observed for KCg-m in the FlyWire connectome (Figure 5G) raises the question of how dimensionality varies with *K* for each of the KC types. Using the neural network rate model of Litwin-Kumar *et al.*^70^, we calculated dimensionality as a function of *K* for each of the KC types, using the observed KC counts, ALPN->KC connectivity, and global inhibition from APL. This analysis revealed that optimal values for *K* are lower for KCg-m in FlyWire than the hemibrain (Figure 5K), similar to the observed values.

Taken together these results demonstrate that for KCg-m the brain compensates for a developmental perturbation by changing a single parameter: the number of odour channels each KC samples from. In contrast, KCa’b’s, which are also more numerous in FlyWire than in the hemibrain, appear to employ a hybrid strategy of reduced *K* combined with a reduction in ALPN→KCa’b’ connection strength (<u>Figure S5F</u>). These findings contradict earlier studies where a global increase in KC numbers through genetic manipulation triggered an increase in ALPN axon boutons (indicating an compensatory increase in excitatory drive to KCs) and a modest increase in KC claws (suggesting an increase rather than decrease in *K*)^73, 74^. This may be due to the differences in the nature and timing of the perturbation in KC cell number, and KC types affected.

### Toward multi-connectome cell typing

As the first dense, large-scale connectome of a fly brain, the hemibrain dataset proposed over 5,000 previously unknown cell types in addition to confirming 512 previously reported types recorded in virtualflybrain.org^2^. Since this defines a *de facto* standard cell typing for large parts of the fly brain, our initial work plan was simply to re-identify hemibrain cell types in FlyWire, providing a critical resource for the fly neuroscience community. While this was successful for 61% of hemibrain cell types (Figure 3), 39% could not yet be validated. Given the great stereotypy generally exhibited by the fly nervous system, this result is both surprising and interesting.

We can imagine two basic categories of explanation. First, that through ever closer inspection we may successfully reidentify these missing cell types. Second, that these definitions, mostly based on a single brain hemisphere, might not be robust to variation across individuals. Distinguishing between these two explanations is not at all straightforward. We began by applying across-dataset connectivity clustering to large groups of unmatched hemibrain and flywire neurons. We observed that most remaining hemibrain types showed complex clustering patterns, that both separated neurons from the same proposed cell type and recombined neurons of different proposed hemibrain types.

While it is always more difficult to prove a negative result, these observations strongly suggest that the majority of the remaining 2,021 hemibrain types are not robust to inter-individual variation. We therefore developed a new definition of cell type that uses inter-animal variability: *A cell type is a group of neurons that are each more similar to a group of neurons in another brain than to any other neuron in the same brain*. This definition can be used with different similarity metrics, but for connectomics data, a similarity measure incorporating morphology and/or connectivity is most useful. Our algorithmic implementation of this definition operates on the co-clustering dendrogram by finding the smallest possible clusters that satisfy two criteria (Figure 6A):

1. Each cluster must contain neurons from all three hemispheres (hemibrain, FlyWire right and FlyWire left).
2. Within each cluster, the number of neurons from each hemisphere must be approximately equal.

Determining how to cut a dendrogram generated by data clustering is a widespread challenge in data science for which there is no single satisfactory solution. A key advantage of the cell type definition we propose is that it provides very strong guidance about how to assign neurons to clusters. This follows naturally from the fact that connectome data provides us with *all* neurons in each dataset, rather than a random subsample. This advantage of completeness is familiar from analogous problems such as the ability to identify orthologous genes when whole genomes are available^75^.

Analysis of the hemibrain cell type AOTU063 provides a relatively straightforward example of our approach (Figure 6B; <u>Figure S7</u>). Morphology-based clustering generates a single group, comprising all four AOTU063 neurons from each of the three hemispheres. However, clustering based on connectivity reveals two discrete groups, with equal numbers of neurons from each hemisphere, suggesting that this type should be split further. Here, algorithmic analysis across multiple connectomes reveals consistent connectivity differences between subsets of AOTU063 neurons.

**Figure 6:**
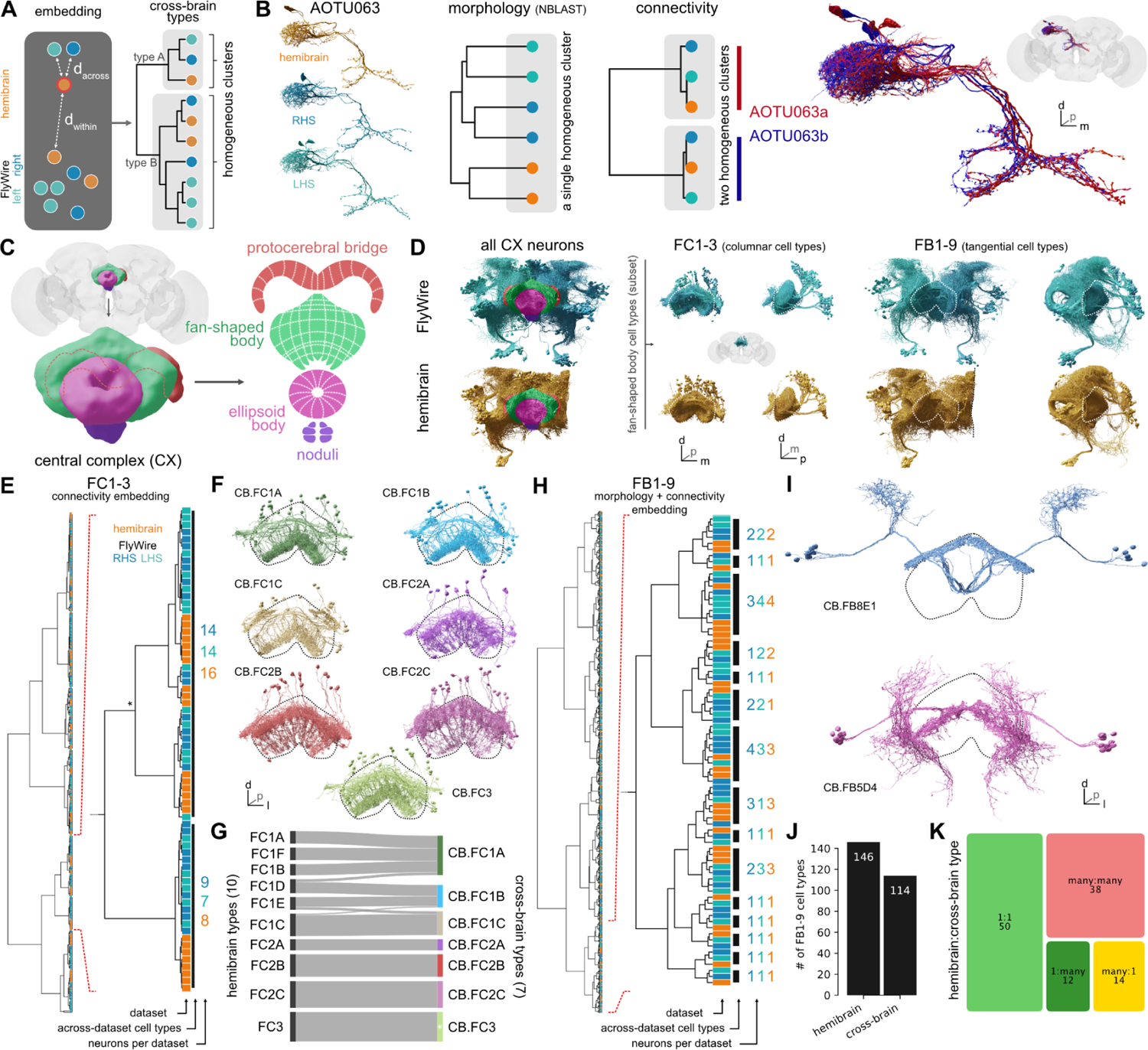
Across-brain cell typing. A. Cell type is defined as a group of neurons that are each more similar to a group in another brain than to any neurons in the same brain. We expect cell type clusters to be homogenous, i.e. contain neurons from all three hemispheres in approximately even numbers. **B** Example of a hemibrain cell type (AOTU063) that is morphologically homogeneous but has two cross-brain consistent connectivity types and can hence be split. **C** Main neuropils making up the central complex (CX). **D** Overview of all CX cells (left) and two subsets of fan-shaped body (FB, dotted outlines) cell types: FC1-3 and FB1-9 (right). **E** Hierarchical clustering from connectivity embedding for FC1-3 cells. Zoom in shows cross-brain cell type clusters. Asterisk marks a cluster that was manually adjusted. **F** Renderings of FC1-3 across-brain types; FB outlined. The tiling of FC1-3 neurons can be discerned. **G** Flow chart comparing FC1-3 hemibrain and cross-brain cell types. Colours correspond to those in F. **H** Hierarchical clustering from combined morphology + connectivity embedding for FB1-9. Zoom in shows cross-brain cell type clusters. **I** Examples from the FB1-9 cross-brain cell typing. Labels are composed from CB.FB{*layer*}{*hemilineage-id*}{*subtype-id*}; FB outlined. See <u>Figure S6</u> for full atlas. **J** Number of hemibrain vs cross-brain FB1-9 cell types. **K** Mappings between hemibrain and cross-brain cell types. See <u>Figure S6</u> for a detailed flow chart.

Is this approach applicable to more challenging sets of neurons? We tested it by setting aside the hemibrain types and carrying out a complete retyping of neurons in the central complex (CX) (Figure 6C), a centre for navigation in the insect brain that has been subject to detailed connectome analysis^61^. We selected two large groups of neurons innervating the fan-shaped body (FB) that show a key difference in organisation. The first group, FC1-3 (357 neurons in total) consists of columnar cell types that tile the FB innervating adjacent non-overlapping columns. The second group, FB1-9 (897 neurons in total), contains tangential neurons where neurons of the same cell type are precisely co-located in space^61^ (Figure 6D). Standard NBLAST similarity assumes that neurons of the same cell type overlap closely in space; while this is true for most central brain types it does not hold for repeated columnar neurons such as those in the optic lobe or these FC neurons of the fan-shaped body. We therefore used a connectivity-only distance metric co-clustering across the three hemispheres. This resulted in 7 FC clusters satisfying the above criteria (Figure 6E,F). Five of these “cross-brain” types have a one-to-one correspondence with hemibrain types, while two are merges of multiple hemibrain types; only a small number of neurons are recombined across types (Figure 6G). For the second group, FB1-9, a combined morphology and connectivity embedding was employed. Co-clustering across the three hemispheres generated 114 cell types compared to 146 cell types in the hemibrain (Figure 6H-J). 44% of these types correspond one-to-one to a hemibrain cell type; 11% are splits (1:many), 12% are merges (many:1), and 33% are recombinations (many:many) of hemibrain cell types (Figure 6K; <u>Figure S6B</u>). The 67% (44+11+12) success rate of this *de novo* approach in identifying hemibrain cell types is slightly higher than the 61% achieved in our directed work in (Figure 3); it is consistent with the notion that further effort could still identify some unmatched hemibrain types, but that the majority will likely require retyping.

All the preceding efforts have focused on cell typing neurons contained by both FlyWire and the hemibrain. But what about the extensive regions of the brain covered only by FlyWire and not by hemibrain? Based on the lessons learned from the joint analysis of hemibrain and FlyWire, a conservative heuristic was used to identify 1,458 cell types based solely on the two hemispheres of FlyWire (<u>Figure S8</u>); these neurons are either completely absent or severely truncated in the hemibrain. We can therefore provide types for all descending neurons (DNs, see <u>Methods</u>) and many brain-spanning neurons (Figure <u>S8E-H</u>). In summary, cell typing based on joint analysis of multiple connectomes proved capable of recapitulating many cell types identified in the hemibrain dataset, while also defining new candidate cell types that are consistent both within and across datasets.

Further validation of the new types proposed by this approach will depend on additional *Drosophila* connectomes, which are forthcoming. We predict that cell types defined this way will be substantially more robust than cell types defined from a single connectome alone.

## Discussion

In this study, we generate human-readable annotations for all neurons in the fly brain at various levels of granularity: superclass, cell class, hemilineage, morphology group, cell type. These annotations provide salient groupings which have already been proven useful not just in our own analyses, but also many of those in our companion paper^1^ as well as other publications in the FlyWire paper package introduced there, and to researchers now using the online platforms Codex (<u>codex.flywire.a</u>i) and FAFB-FlyWire CATMAID spaces (<u>fafb-flywire.catmaid.org</u>). Hemilineage annotations also provide a key starting point to link the molecular basis of the development of the central brain to the wiring revealed by the connectome; such work has already begun in the more repetitive circuits of the optic lobe^76^.

The cell type atlas that we provide of 4,552 cell types is not the largest ever proposed (the hemibrain had 5,235), and very new work in the mouse brain has offered up to 5,000 molecular clusters^77–79)^. However it is, by some margin, the largest ever *validated* collection of cell types.^34^ In *C. elegans*, the 118 cell types inferred from the original connectome have been clearly supported by analysis of subsequent connectomes and molecular data^4, 5, 80–82^. In a few cases in mammals, it has been possible to produce catalogues of order 100 cell types that have been validated by multimodal data e.g. in the retina or motor cortex^35, 83, 84^. While large scale molecular atlases in the mouse produce highly informative hierarchies of up to 5,000 clusters ^77–79^, they do not yet try to define terminal cell types – the finest unit that is robust across individuals – with precision. Here, to our knowledge for the first time, we test over 5,000 predicted cell types, resulting in 3,094 validated cell types using 3 hemispheres of connectome data. Informed by this we use the FlyWire dataset to propose an additional 1,458 cell types.

### Lessons for cell typing

Our experience of cell typing the FlyWire dataset together with our earlier participation in the hemibrain cell typing effort leads us to draw a number of lessons. First, we think that it is helpful to frame cell types generated in one dataset as predictions or hypotheses that can be tested either through additional connectome data or data from other modalities. Related to this, although the two hemispheres of the same brain can be treated as two largely independent datasets, we do see evidence that variability can be correlated across hemispheres (Figure 4). Therefore we recommend the use of three or more hemispheres to define and validate new cell types both because of increased statistical confidence and because across-brain comparisons are a strong test of cell type robustness. Third, there is no free lunch in the classic lumping versus splitting debate. The hemibrain cell typing effort preferred to split rather than lump cell types, reasoning that over-splitting could easily be remedied by merging cell types at a later date^3^. Although this approach seemed reasonable at the time, it appears to have led to cell types being recombined: when using a single dataset even domain experts may find it very hard to distinguish conserved differences between cell types from inter-individual noise. Fourth, although some recent studies have argued that cell types are better defined by connectivity than morphology, we find that there is a place for both. For example when faced with the task of aligning two connectomes, we recommend starting by using morphology to identify “landmark” cell types. These shared cell type labels can then be used to define connection similarity for neurons in the two datasets. Fifth, and related to this, we find that across-dataset connection similarity is an extremely powerful way to identify cell types. However, connectivity-based typing is typically used recursively and especially when used within a single dataset this may lead to selection of idiosyncratic features. Moreover neurons can connect similarly but come from a different developmental lineage, or express a different neurotransmitter, precluding them from sharing a cell type. Combining these two points, we would summarise that matching by morphology appears both more robust and sometimes less precise, whereas connectivity matching is a powerful tool that must be wielded with care.

In conclusion, connectome data is particularly suitable for cell typing: it is inherently multimodal (by providing morphology and connectivity) while the ability to see all cells within a brain (completeness) is uniquely powerful. Our multi-connectome typing approach (Figure 6) provides a robust and efficient way to use such data; cell types that have passed the rigorous test of across-connectome consistency are very unlikely to be revised (permanence). We suspect that connectome data will become the gold standard for cell typing. Linking molecular and connectomics cell types will therefore be key. One promising new approach is exemplified by the prediction of neurotransmitter identity directly from EM images^38^ but many others will be necessary. In the introduction we set out to answer three questions, which we will now discuss in closing.

### Can we simplify the connectome graph to aid automated or human analysis?

Cell typing reduces the complexity of the connectome graph. This has important implications for analysis, modelling, experimental work and developmental biology. For example, using the cell types identified so far, we can reduce 45,675 nodes in the raw connectome graph into a cell type graph with 4,552 nodes; the number of edges is similarly reduced. This should significantly aid human reasoning about the connectome. It will also make numerous network analyses possible as well as substantially reduce the degrees of freedom in brain scale modelling^85, 86^. It is important to note that while collapsing multiple cells for a given cell type into a single node is often desirable, other use cases such as modelling studies may still need to retain each individual cell. However, if key parameters are determined on a per cell type basis, then the complexity of the resultant model can be much reduced. Recently Lappalainen et al.^85^ optimised and analysed a highly successful model of large parts of the fly visual system with just 734 free parameters by using connectomic cell types.

For *Drosophila* experimentalists using the connectome, cell typing identifies groups of cells that likely form functional units. Most of these are linked though <u>virtualflybrain.org</u> to the published literature and in many cases to molecular reagents. Others will be more easily identified for targeted labelling and manipulation after typing. Finally, cell typing effectively compresses the connectome, reducing the bits required to store and specify the graph. For a fly-sized connectome this is no longer that important for computerised analysis, but it may be important for development. Zador^87^ has argued that evolution has selected highly structured brain connectivity enabling animals to learn very rapidly, but that these wiring diagrams are far too complex to be specified explicitly in the genome; rather they must be compressed through a “genomic bottleneck” which may itself have be a crucial part of evolving robust and efficient nervous systems. If we accept this argument, lossy compression based on aggregating nodes with similar cell type labels, approximately specifying strong edges and largely ignoring weak edges would reduce the storage requirements by orders of magnitude and could be a specific implementation of this bottleneck.

### How do we know which edges are important?

The question of which of the 16.5M edges in the connectome to pay attention to is critical for its interpretation. Intuitively we assume that the more synapses are found to connect two neurons, the more important that connection must be. There is some very limited evidence in support of this assumption correlating anatomical and functional connectivity^88, 89^ (compare in mammals^90^). In lieu of physiological data, we postulate that edges critical to brain function should be consistently found across brains. By comparing connections between cell types identified in 3 hemispheres we find that edges stronger than 10 synapses or 1.1% of the target’s inputs have a greater than 90% chance to be preserved (Figure 4F). This provides a simple heuristic for determining which edges are likely to be functionally relevant. It is also remarkably consistent with new findings from the larval connectome where left-right asymmetries in connectivity vanish after removing edges weaker than <1.25%^91^. It is important to note though that edges falling below the threshold might still significantly contribute to the brain’s function.

We further address an issue that has received little attention (but see^92^): the impact of technical factors (such as segmentation, proofreading, synapse detection) and biological variability on the final connectome and how to compensate for it. In our hands, a model of technical noise could explain up to 30% difference in edge weights. While this model was made specifically for the two hemispheres of FlyWire, it highlights the general point that a firm understanding of all sources of variability will be vital for the young field of comparative connectomics to distinguish real and artificial differences.

### Have we collected a snowflake?

The field of connectomics has long been criticised for unavoidably low N^93, 94^: is the brain of a single specimen representative for all? For insects, there is a large body of evidence for morphological and functional stereotypy, although this information is only available for a minority of neurons and much less is known about stereotyped connectivity^34, 95, 96^. For vertebrate brains the situation is less clear again; it is generally assumed that subcortical regions will be more stereotyped, but cortex also has conserved canonical microcircuits^97^ and recent evidence has shown that some cortical elements can be genetically and functionally stereotyped^98–100^. Given how critical stereotypy is for connectomics it is important to check whether that premise actually still holds true at synaptic resolution.

For the fly connectome, the answer to our question is actually both more nuanced and more interesting than we initially imagined. Based on conservation of edges between FlyWire and hemibrain hemispheres over 50% of the connectome graph is a snowflake! Of course these non-reproducible edges are mostly weak. Our criterion for strong (highly reliable) edges retains between 7-16% of edges and 50-70% of synapses.

We previously showed that the early olfactory system of the fly is highly stereotyped in both neuronal number and connectivity^60^. That study used the same EM datasets - FAFB and the hemibrain - but was limited in scope since only manual reconstruction in FAFB was then available. We now analyse brain-wide data from two brains (FlyWire and the hemibrain) and three hemispheres to address this question and find a high degree of stereotypy at every level: neuron counts are highly consistent between brains, as are connections above a certain weight. However, when examining so many neurons in a brain, we can see that cell counts are very different for some neurons; furthermore neurons occasionally do something unexpected (take a different route or make an extra branch on one side of the brain). In fact, we hypothesise that such stochastic differences are unnoticed variability present in most brains; this is reminiscent of the observation that most humans carry multiple significant genetic mutations. We did observe one example of a substantial biological difference that was consistent across hemispheres but not brains: the number of the KCg-m neurons in the mushroom bodies is almost twice as numerous in FlyWire than in the hemibrain. Intriguingly, we found evidence that the brain compensates for this perturbation by modifying connectivity (Figure 5).

In conclusion, we can say that we have *not* collected a snowflake; the core FlyWire connectome is highly conserved and the accompanying annotations will be broadly useful across all studies of *Drosophila melanogaster*. However, our analyses show the importance of calibrating our understanding of biological (and technical) variability – as has recently been done across animals in *C. elegans*^82^ and within hemispheres in larval *Drosophila*^91, 101^. This will be crucial to identifying true biological differences in sexually dimorphic circuits or changes due to learning using future connectomes.

## Supplemental Figures

**Supplemental Figure S1.**
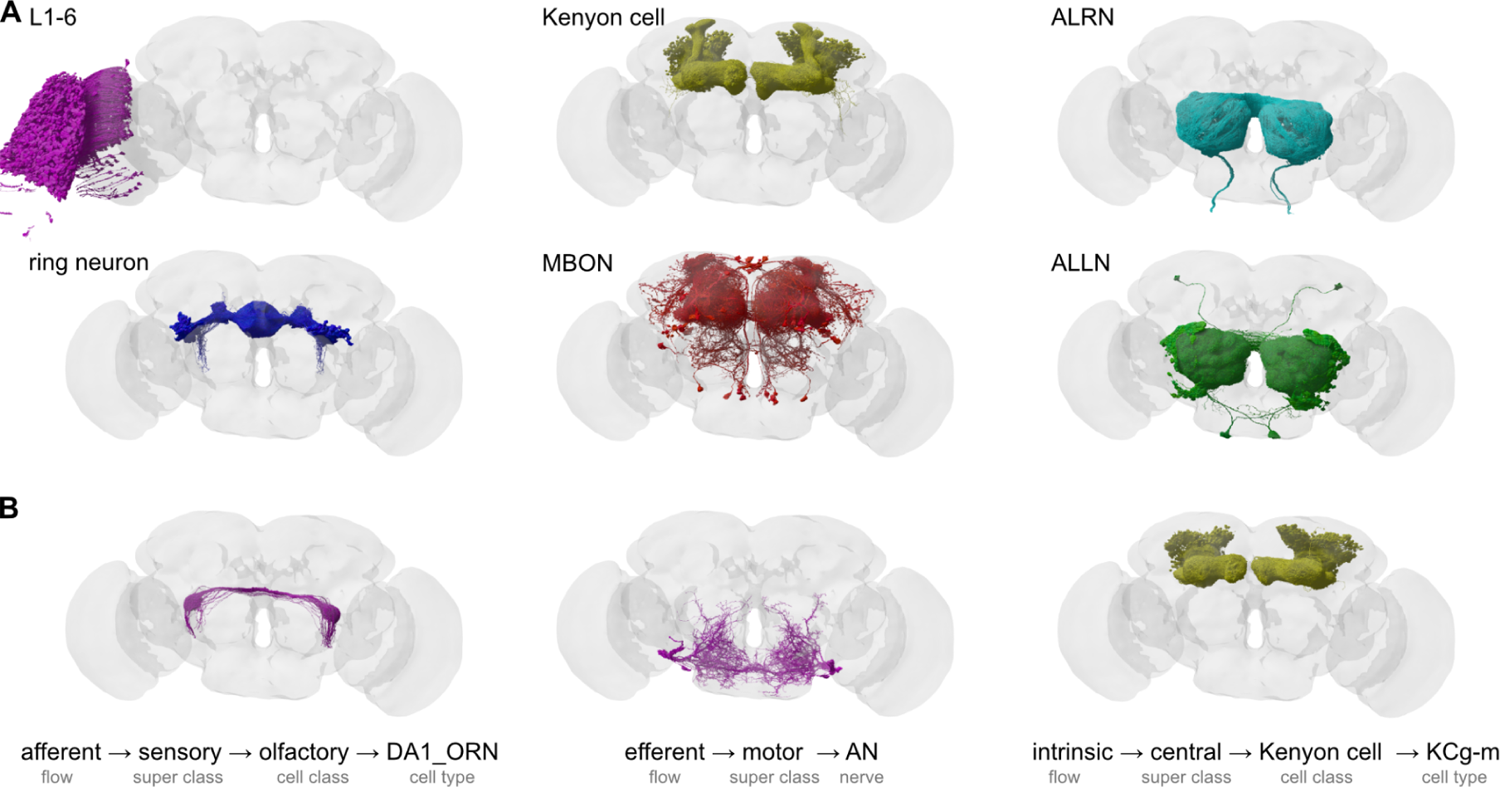
A Examples for cell class annotations. **B** Examples for labels derived from the hierarchical annotations. Abbreviations: ALRN, antennal lobe receptor neuron; MBON, mushroom body output neuron; ALLN, antennal lobe local neuron; ORN, olfactory receptor neuron; AN, antennal nerve.

**Supplemental Figure S2.1.**
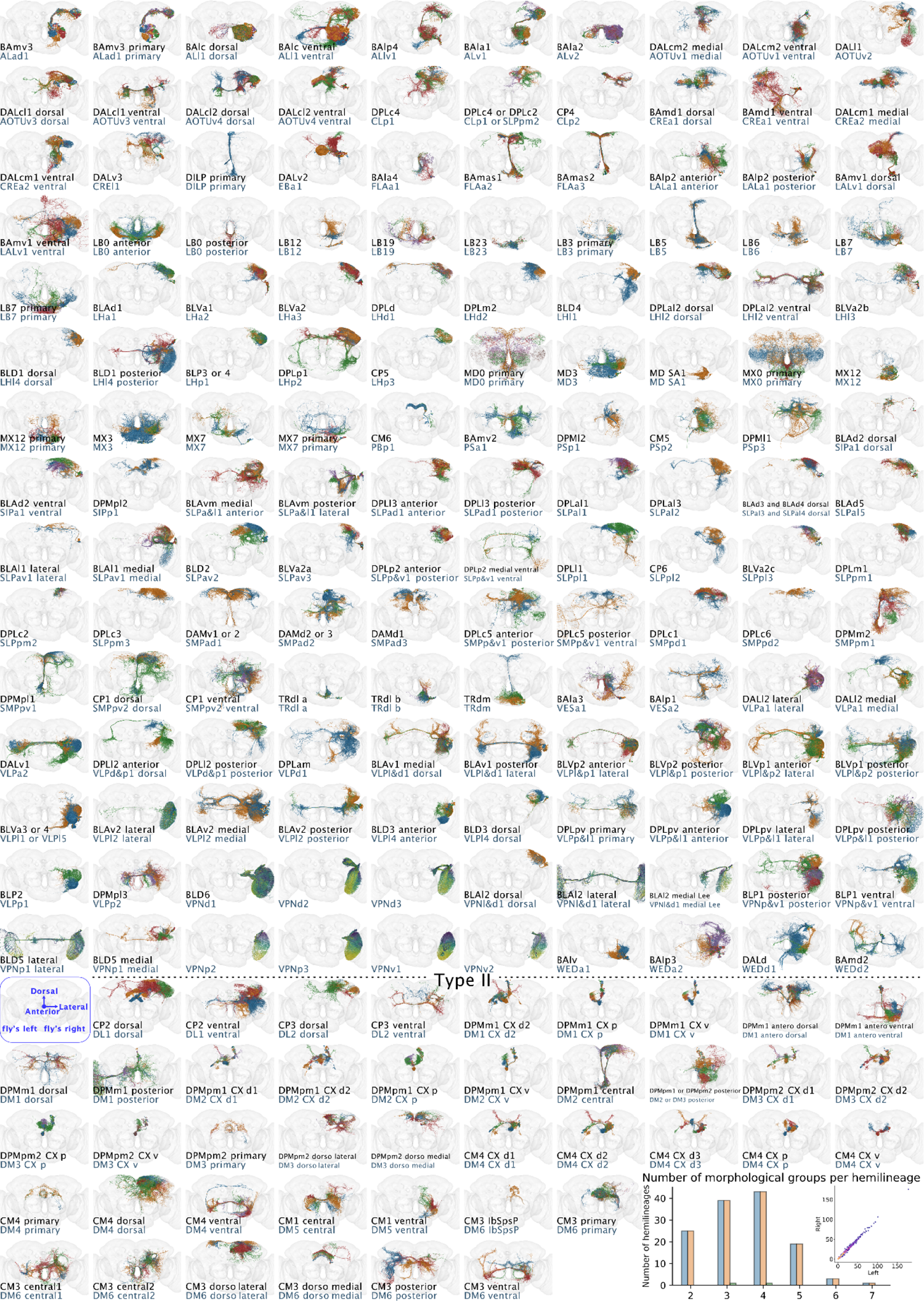
Anterior views of neurons within a hemilineage (based on^53, 102^), or neurons whose cell bodies form a cluster in a lineage clone (also referred to as “hemilineages’’ hereafter), based on the light-level data from^46–49, 103^. The names of the hemilineages are at the bottom of each panel (top: Hartenstein nomenclature; bottom: ItoLee nomenclature). The snapshots only include neurons with cell bodies on the right hemisphere, and the central unpaired lineages. Except for the hemilineages that tile the optic lobe, the neurons are coloured by morphological groups, obtained by the ‘elbow method’ (Methods, <u>Hemilineage</u> <u>annotations section</u>). The neurons that form cohesive tracts with their cell body fibres in the Type II lineages (see Methods) are at the lower part of the panels. The first panel of the “Type II’ section is for orientation purposes. The bottom right panel is a histogram of the number of morphological groups per hemilineage (blue: left; yellow: right; green: centre). Inset is the number of neurons per hemisphere for each morphological group, with points coloured by their density (yellow: denser). Corresponding group names, together with FlyWire and neuroglancer links are available in <u>Supplementary Files 2 and 3</u>.

**Supplemental Figure S2.2.**
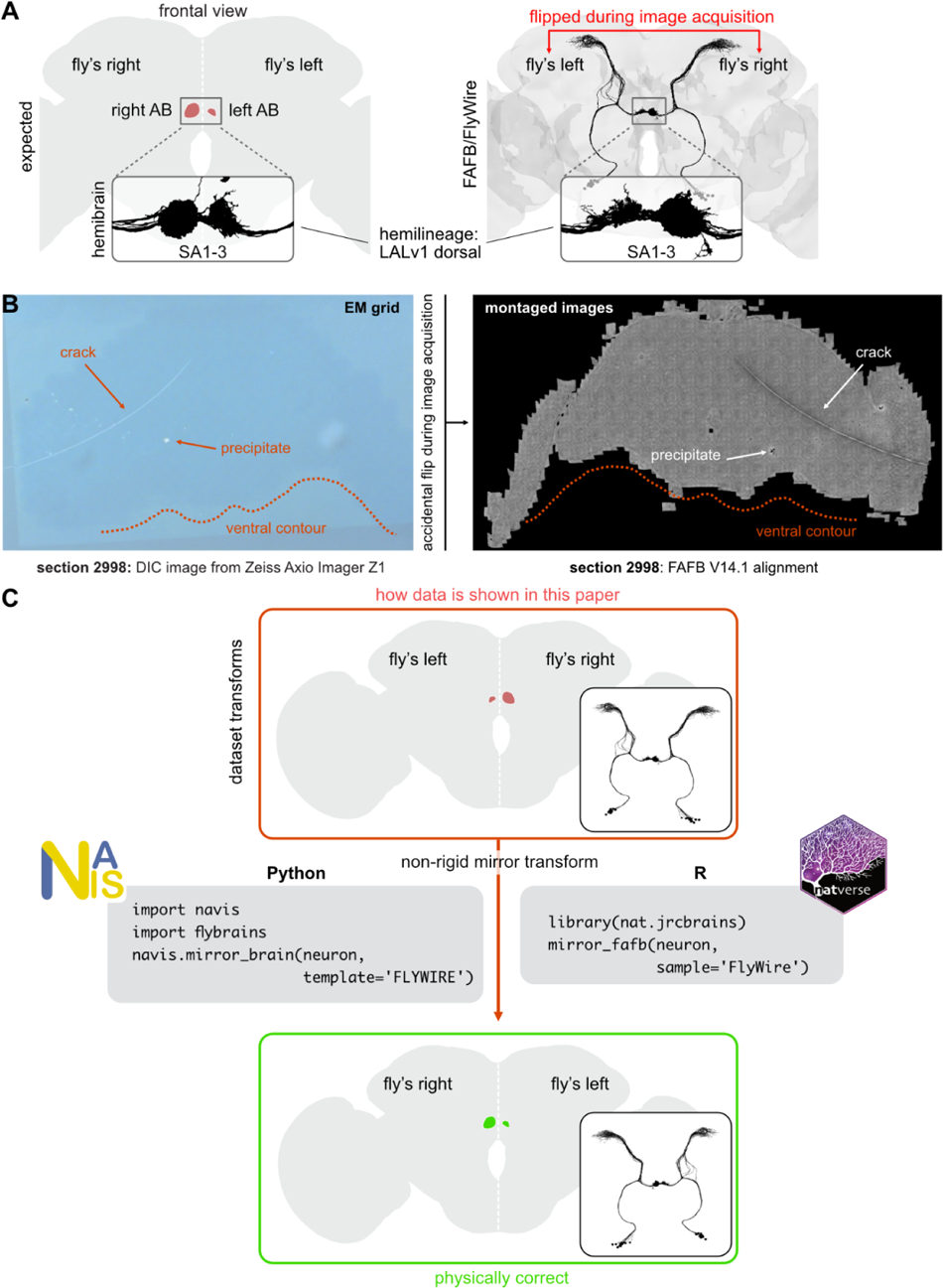
**A** The asymmetric body (AB) is expected to be larger on the fly’s right. The adult fly brain is conventionally shown in frontal views in 2D projections; this would place the fly’s right on the left of the page. In FAFB/FlyWire the situation initially appeared inverted. Insets show axons of SA1-3 neurons which form the major input to the AB. **B** Image of a brain section on the original EM grid (left) and the final image montage as shown in neuroglancer/CATMAID (right). Various landmarks are shown to illustrate the flip along the x-axis that explains the inversion. **C** Showcase of how to correct the inversion of FAFB/FlyWire data. For technical reasons, it was not possible to flip the whole FAFB volume and associated data. Therefore this must be corrected post hoc. Code samples show how this can be done for e.g. mesh or skeleton data using Python or R.

**Supplemental Figure S3.**
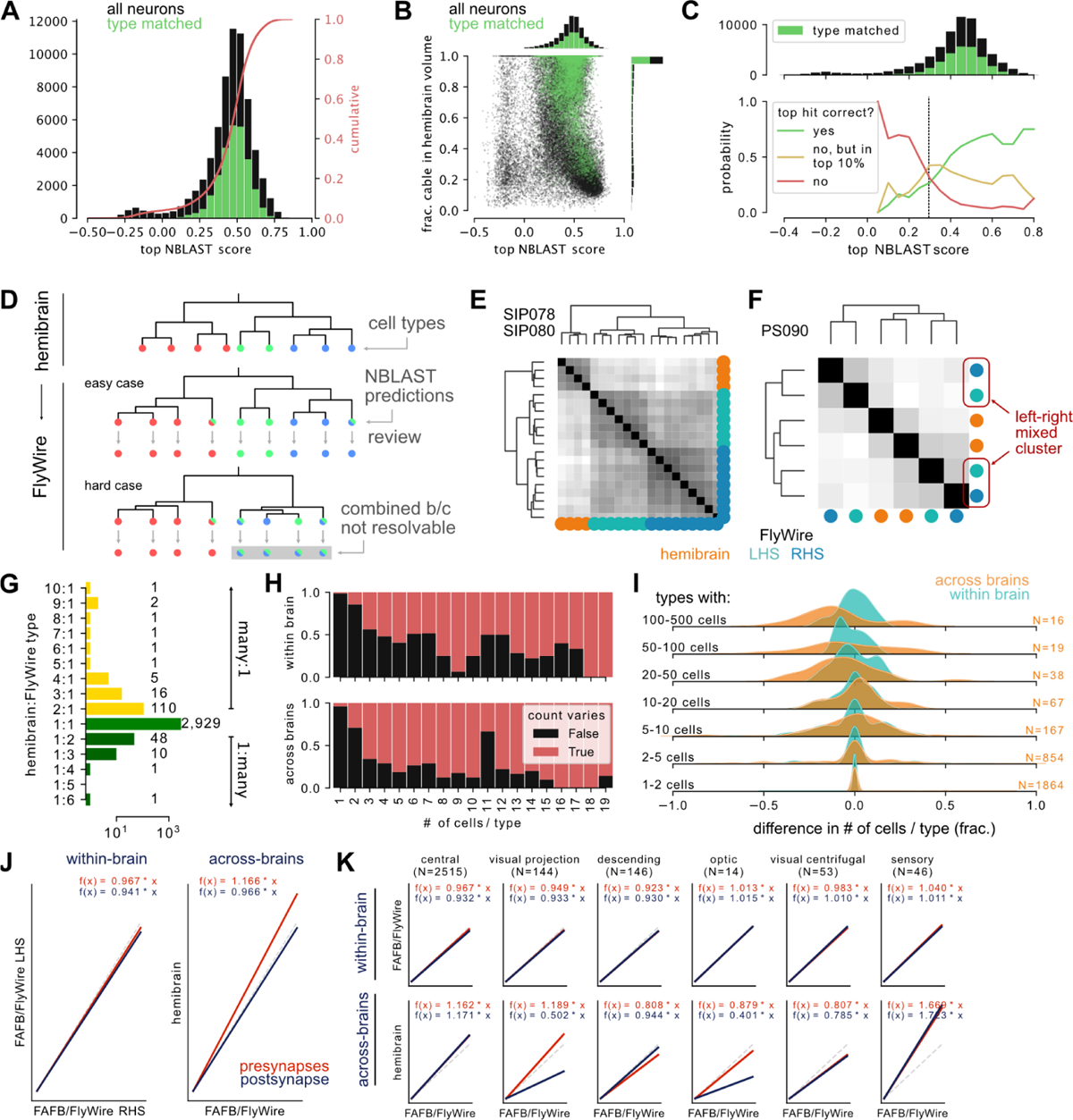
**A** Distribution of top FlyWire → hemibrain NBLAST scores. **B** Top NBLAST scores vs fraction of neuron contained within hemibrain volume. Heavily truncated neurons typically produce bad scores. **C** Top: distribution of top NBLAST scores and fraction which was type matched. Bottom: probability that the correct hit was the top NBLAST hit (green) or at least among (yellow) the top 10% (by score) as a function of the top NBLAST score. **D** Explanation of match review process: where possible the within-dataset morphological clustering was taken into account. **E,F** Cross-brain co-clustering of the NBLAST scores for example cell types in Figure 3. **G** Counts for 1:many and many:1 type matches. These also include types derived from previously untyped hemibrain neurons. **H** Fraction of cell types showing a difference in cell counts within (left/right, top) and across (bottom) brains. **I** Distribution of cell count differences. **J** Robust linear regression (Huber w/ intercept at 0) for within- and across-dataset pre/postsynapse counts from Figure 3H. **K** Same data as in J but separated by superclass. Slopes are generally close to 1: 1.021 (pre-) and 1.035 (postsynapses, i.e. inputs) between the left and right hemisphere of FlyWire, and 1.176 (presynapses, i.e. outputs) 0.983 (post) between FlyWire and the hemibrain. Note that correlation and slope are noticeably worse for cell types known to be truncated such as visual projection neurons which suggests that we did not fully compensate for the hemibrain’s truncation and that the actual across-brain correlation might be even better.

**Supplemental Figure S4.**
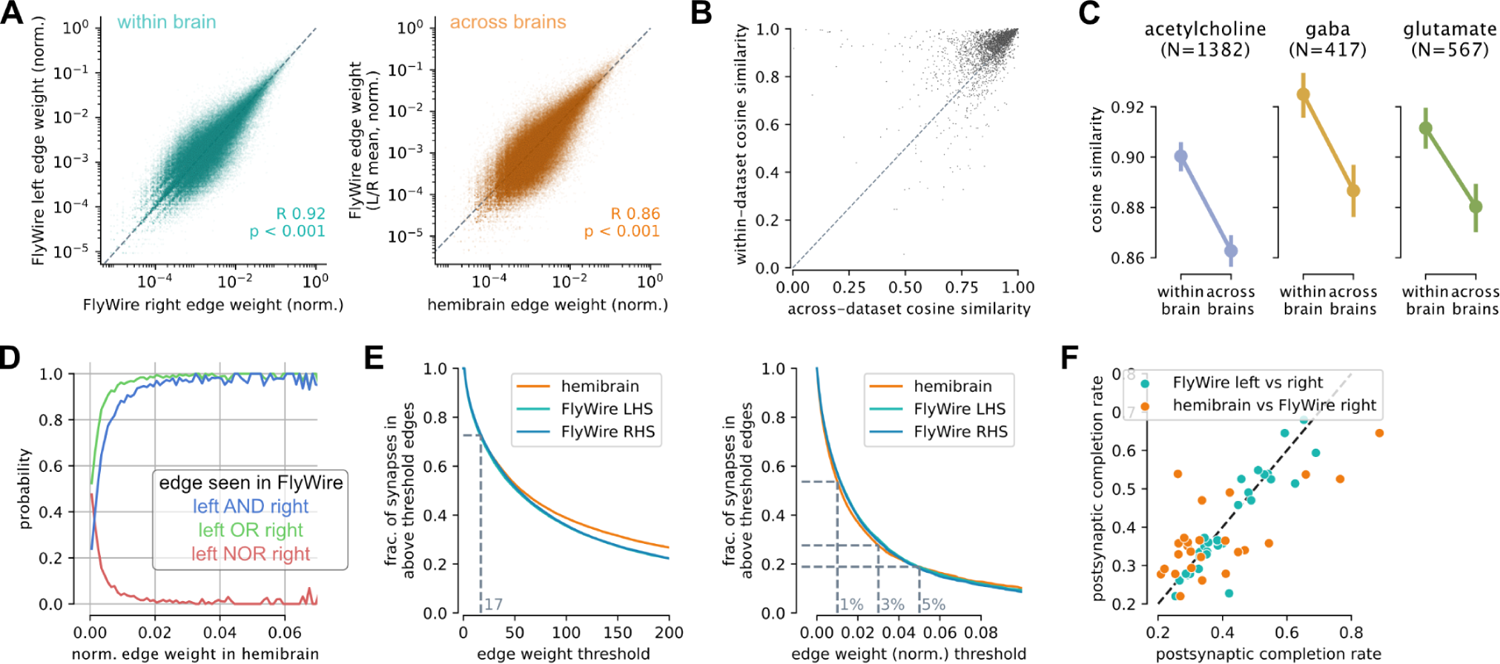
**A** Comparison of normalised edge weights within (left) and across (right) brains. **B** Connectivity cosine connectivity similarity within and across brains. Each datapoint is a cell type identified across the three hemispheres. Size correlates with the number of cells per type. **C** Connectivity cosine similarity separated by neurotransmitter. **D** Probability that an edge present in the hemibrain is found in one, both or neither of the FlyWire hemispheres. **E** Fraction of synapses contained in edges above given absolute (left) and normalised (right) weight. **F** Postsynaptic completion rates. Each datapoint is a neuropil.

**Supplemental Figure S5.**
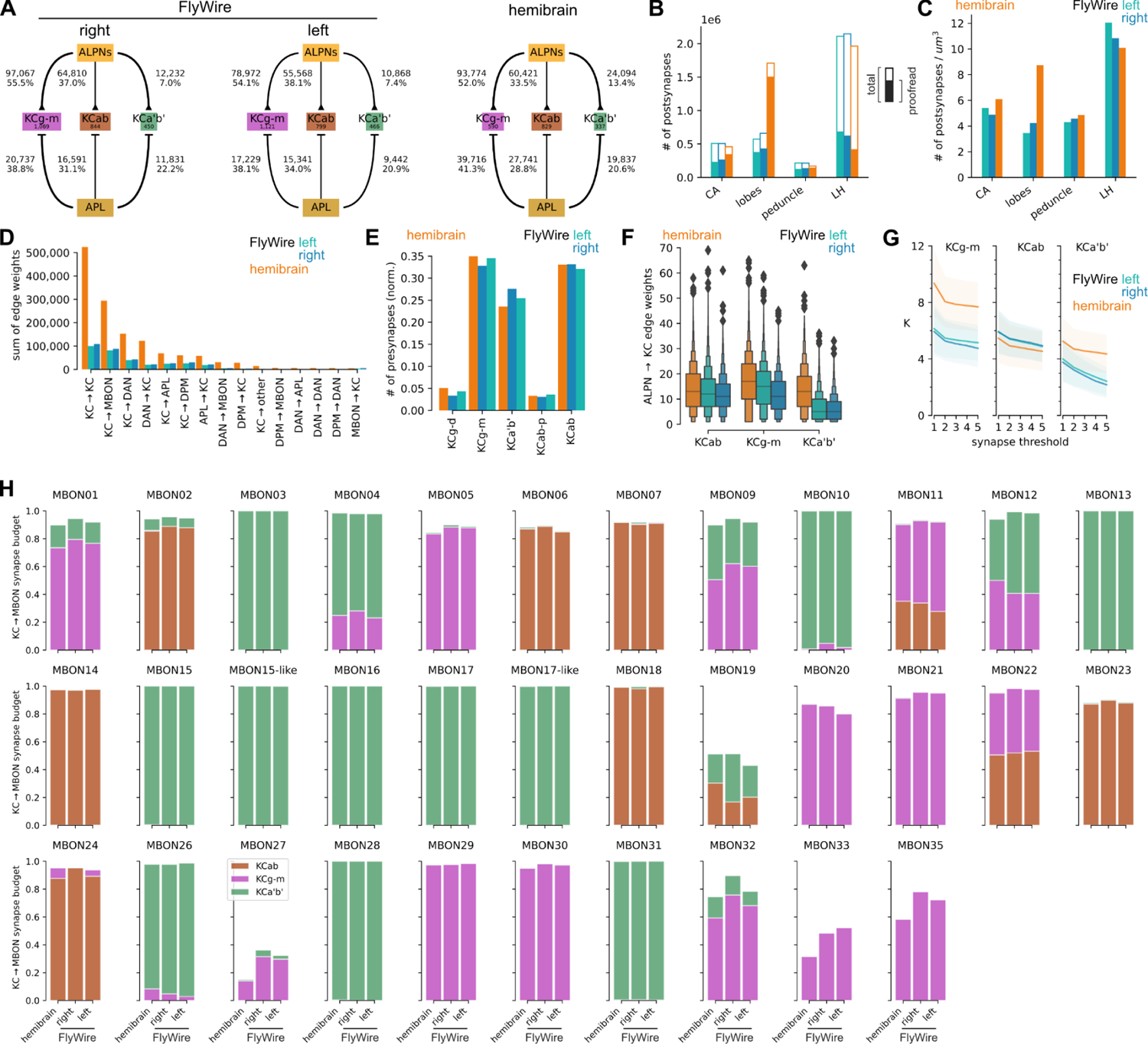
**A** Graph showing ALPN/APL→KC connectivity across the three datasets. Edge labels provide weights both as total synapse counts and normalised to the total output budget of the source. In FlyWire, the mushroom bodies (MB) have 57.2% (left) and 60.7% (right) postsynaptic completion rate while the hemibrain MB has been proofread to 81.3% (see also B). To compensate for this we typically used normalised synapse counts and edge weights. Note that KCab act as an internal control as their numbers are consistent across all hemispheres and we don’t expect to see any changes in their connectivity. **B** Total versus proofread postsynapse counts across MB compartments. Lateral horn (LH) shown for comparison. **C** Postsynapse density across MB compartments. **D** Sums of edge weights between different MB cell classes. **E** Presynapse counts per KC type normalised to the total number of KC synapses per dataset. **F** ALPN→KC edge weights. **G** *K* (# of ALPN types providing input to a single KC) under different synapse thresholds. **H** Fraction of MBON input budget coming from individual KCab, KCg-m and KCa’b’. Abbreviations: CA, calyx; DAN, dopaminergic neuron; ALPN, antennal lobe projection neuron; KC, Kenyon Cell; MBON, mushroom body output neurons.

**Supplemental Figure S6.**
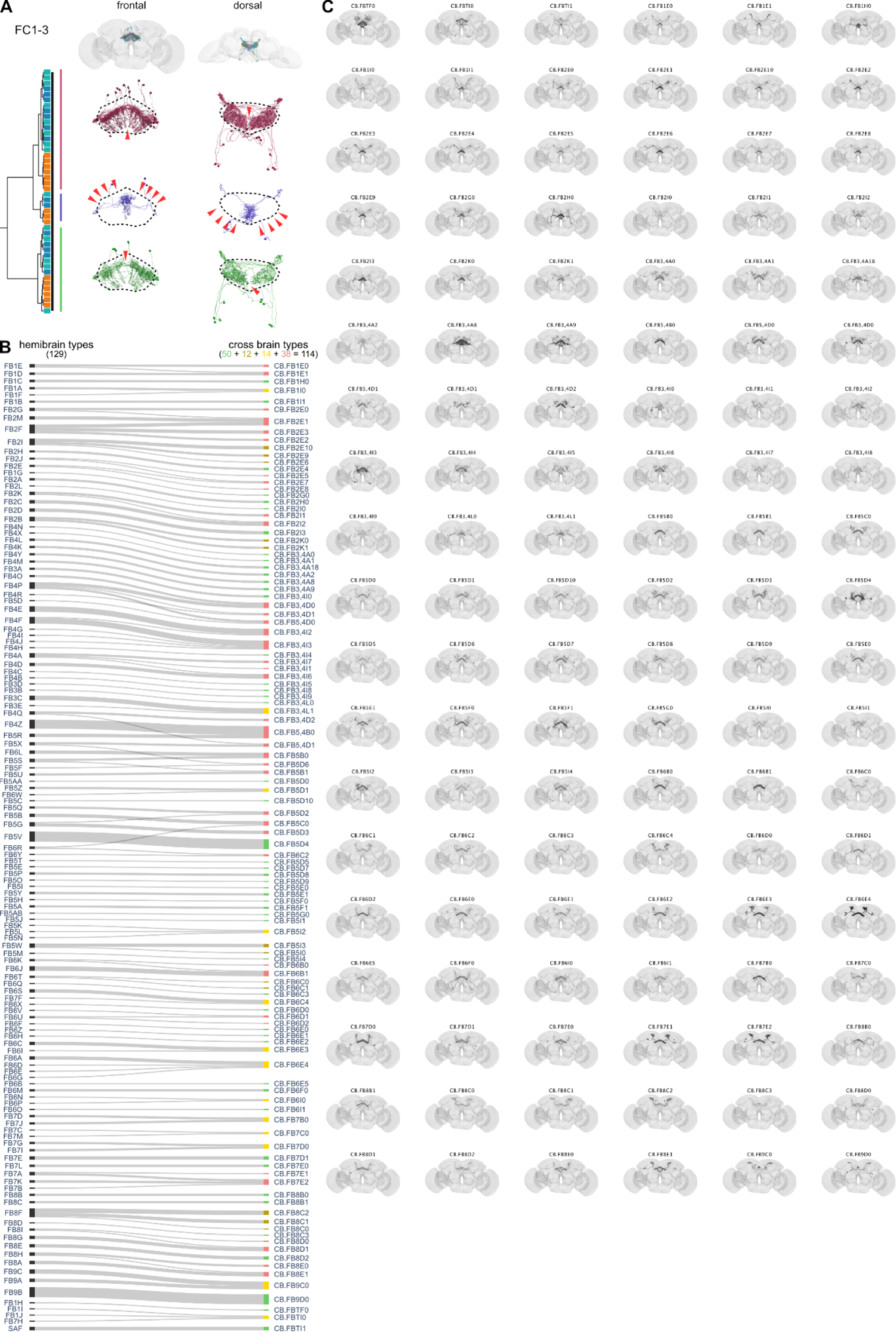
**A** FC1-3 across-brain cluster from Figure 6D (asterisk) that was manually adjusted. This group consists of three sub-clusters that technically fulfil our definition of cell type. They were merged, however, because they individually omit columns of the fan-shaped body (arrowheads) and are complementary to each other. **B** Flow chart comparing FB1-9 hemibrain and cross-brain cell types. Colours correspond to 1:1, 1:many, many:1 and many:many mappings between hemibrain and cross-brain cell types. **C** Renderings of all FB1-9 cross-brain cell types.

**Supplemental Figure S7.**
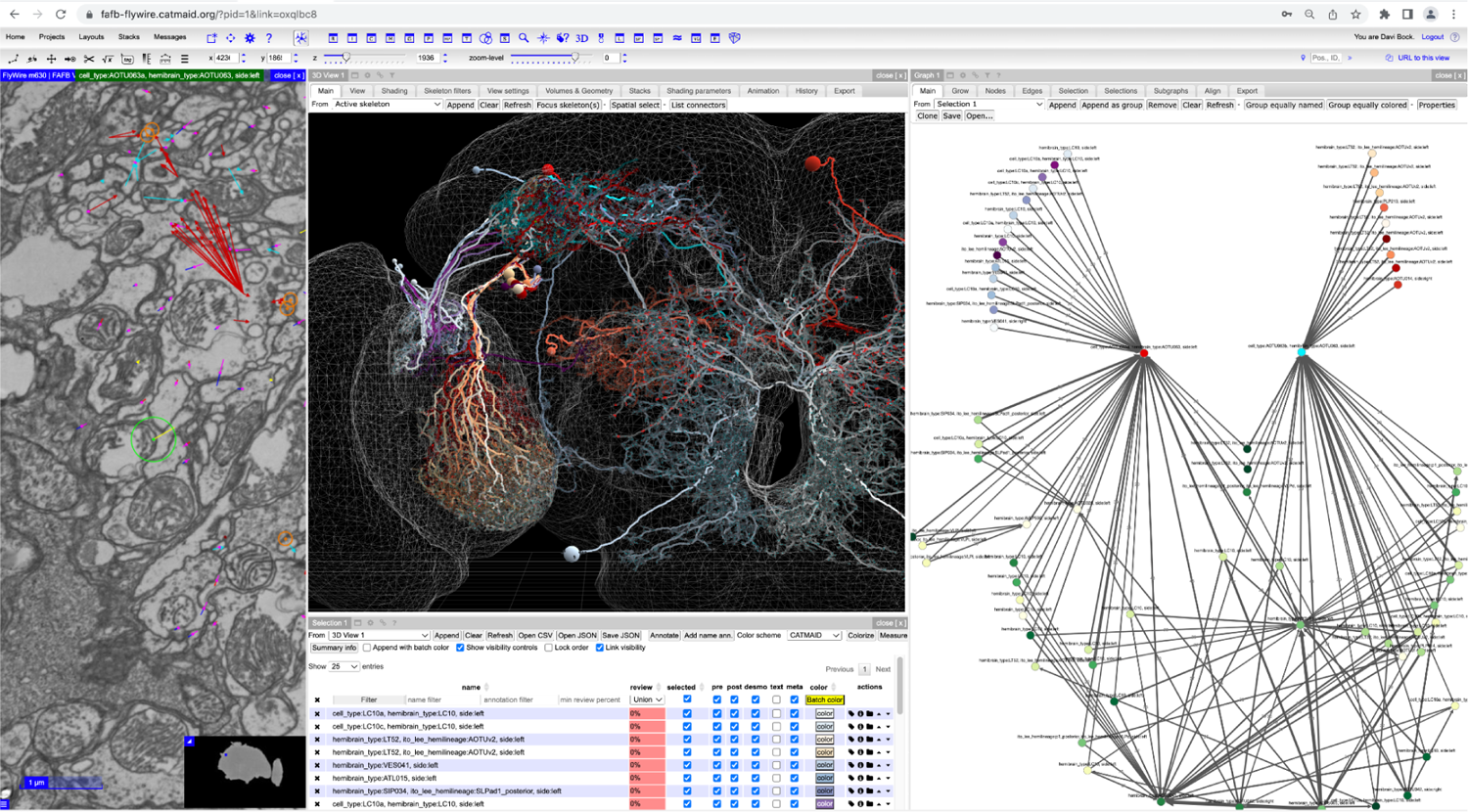
Screenshot demonstrating the use of CATMAID Spaces (https://fafb-flywire.catmaid.org/) to interrogate the FlyWire connectome. Differential inputs to AOTU63a and b are visualised (red and cyan, respectively). The Graph widget was used to show all neurons making 20 or more synapses onto AOTU63a and b, and to show only >=20 synapse connections between these neurons. Neurons whose *only* >=20 synapse connection was to either AOTU63a or b (but not both) were differentially coloured (blue-purples and greens, respectively).

**Supplemental Figure S8:**
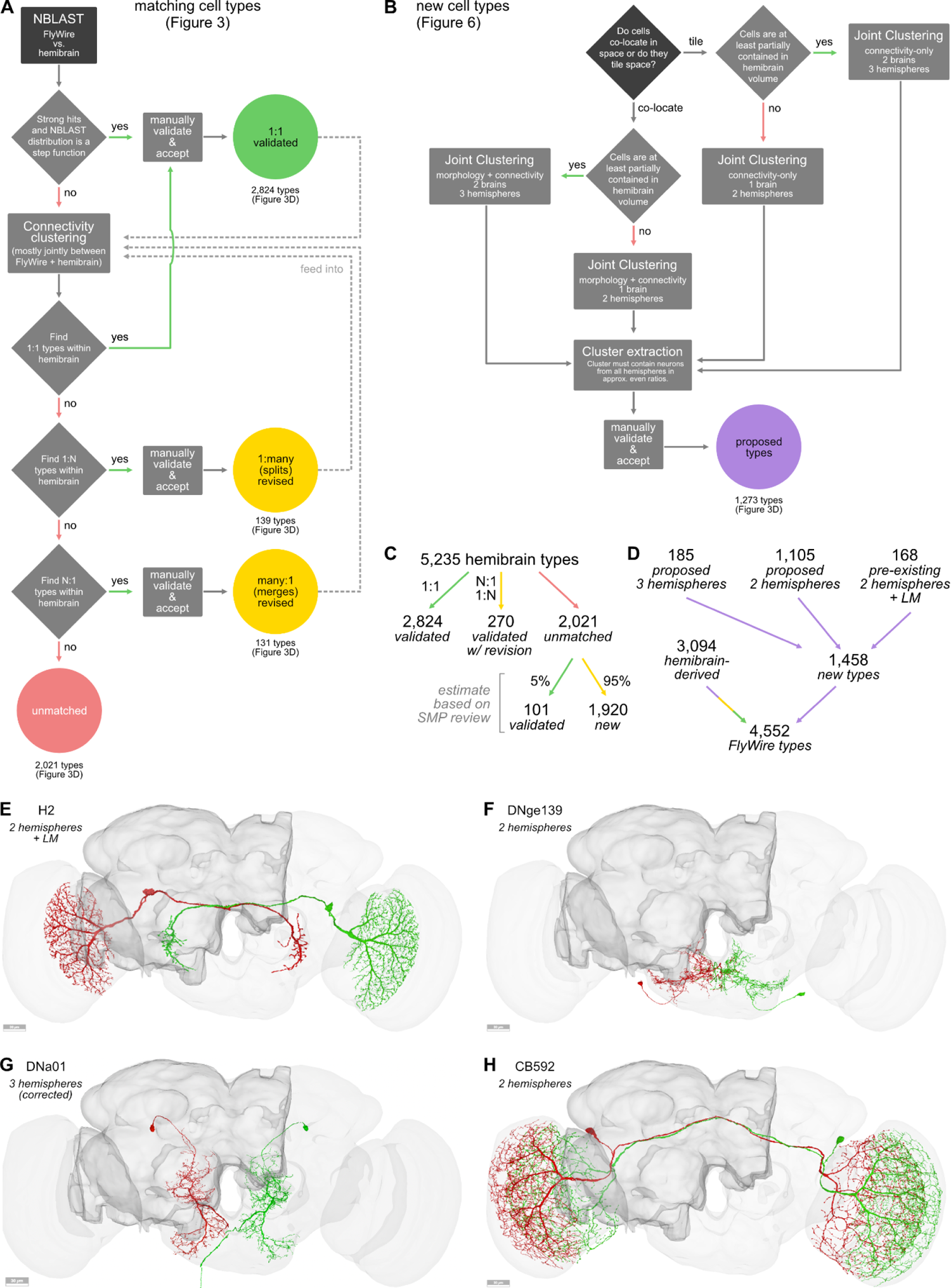
**A** Workflow for matching hemibrain types to FlyWire neurons. **B** Workflow for extracting *de novo* cell types from connectomics data. **C** Summary counts of cell types resulting from workflow (Panel A) to map the 5,235 hemibrain morphological types onto the FlyWire connectome. Based on sampling a subset of types in the superior medial protocerebrum (SMP) we estimate that around 5% of the unmatched hemibrain types could likely be identified in FlyWire using joint morphology-connectivity clustering across connectomes, while 95% will require retyping (data not shown). **D** Breakdown of the 4,552 total cell types provided through our annotations. 3,094 types are based on hemibrain data through the typing workflow (Panel A; 3,094 types). The 1,458 new types are principally based on left vs. right matching within FlyWire, with a small number additionally informed by previously published light microscopy (LM) data (168) and/or large untyped fragments of hemibrain neurons (185). **E-H** Examples of new cell types. H2 is based on left vs right FlyWire clustering plus existing LM data; DNge139 and CB592 are based solely on left vs right FlyWire clustering; DNa01 is based on three hemispheres worth of data but was miss-identified as “VES006” in the hemibrain.

## Supplemental Tables

**Supplementary Table S1:** Example annotations.

| flow | superclass | class | sub class | type | side | hemilineage | neurotransmitter |
| --- | --- | --- | --- | --- | --- | --- | --- |
| afferent | sensory | olfactory | ALRN | ORN_DA1 | left | ALI1_ventral | acetylcholine |
| intrinsic | central | DAN |  | PAM08 | left | CREa1_dorsal | dopamine |
| efferent | descending |  |  | DNa02 | right | WEDd1 | acetylcholine |
| **Supplementary Table S1:** Example annotations. |  |  |  |  |  |  |  |

**Supplementary Table S2:** Glossary for terms used in the top-most layers of the annotation hierarchy. Ontology ID refers to the Virtual Fly Brain database.

| Field | Value | Ontology ID | Definition |
| --- | --- | --- | --- |
| flow | intrinsic |  | Neurons fully contained within the brain. |
|  | afferent |  | Neurons that enter the brain from the periphery or the ventral nerve cord (VNC). |
|  | efferent |  | Neurons that leave the brain towards the periphery or the VNC. |
| superclass | central | FBbt_00059245 | Neurons fully contained within the central brain. |
|  | ascending | FBbt_00048301 | Neurons entering the brain from the VNC. These can be sensory or interneurons. |
|  | descending | FBbt_00047511 | Neurons with soma in the brain that exit the brain towards the VNC. |
|  | endocrine | FBbt_00059246 | Neurons that exit the brain via the NCC towards the retrocerebral complex (corpora cardiaca and corpora allata). |
|  | motor | FBbt_00005123 | Neurons that exit the brain towards the periphery (and are hence assumed to be motor neurons). |
|  | optic | FBbt_00007577 | Neurons fully contained within the optic lobes or the ocellar ganglion. Includes some bilateral neurons (see class field). |
|  | visual projection | FBbt_00048287 | Neurons that have dendrites in the optic lobes or the ocellar ganglion and axons in the central brain. |
|  | visual centrifugal | FBbt_00059244 | Neurons that have dendrites in the central brain and axons in the optic lobes or the ocellar ganglion. |
|  | sensory | FBbt_00005124 | Neurons that enter the brain from the periphery. Note that "ascending" also includes some sensory neurons that we are unable to distinguish from ascending interneurons with any certainty. |

**Supplementary Table S3:** Selection of cell *classes*.

| Class | Subset of superclass | Definition |
| --- | --- | --- |
| bilateral | optic | Optic lobe neurons with projections into the contralateral optic lobes. Sometimes also leave synapses in the central brain. |
| ocellar | visual centrifugal | Projection neurons with dendrites in the brain and axons in the ocellar ganglia. |
|  | visual projection | Projection neurons with dendrites in the ocellar ganglia and axons in the brain. |
|  | optic | Neurons intrinsic to the ocellar ganglia. Cell bodies can be inside the brain though. |
| ALIN | central | “Antennal lobe input neurons”: neurons with dendrites out- and axons inside the antennal lobes. Does not include sensory neurons. |
| ALPN | central | “Antennal lobe projection neurons”: neurons with dendrites in the antennal lobe and axonal projections into the protocerebrum, lateral horn or calyx. |
| ALON | central | “Antennal lobe output neurons”: neurons with dendrites in the antennal lobes and axonal projections outside of the brain that are not canonical ALPNs. |
| ALLN | central | “Antennal lobe local neurons”: neurons with both dendrites and axons contained to the antennal lobes. Can be bilateral. |
| CX | central | Central complex neurons as defined by Hulse *et al.*<sup>61</sup> |
| DAN | central | Dopaminergic neurons (PAM and PPL) whose axons target the mushroom body lobes. |
| Kenyon Cell | central | Neurons with dendrites in the mushroom body calyx whose axons form the parallel fibre system of the mushroom body lobes. |
| MBIN | central | “Mushroom body input neuron”: APL or DPM. |
| LHCENT | central | “Lateral horn centrifugal neurons”: neurons with dendrites in the protocerebrum and axons in the lateral horn. |
| LHLN | central | “Lateral horn local neurons”: neurons with both dendrites and axons contained to the lateral horn. |
| **Supplementary Table S3:** Selection of cell *classes*. |  |  |

### Supplemental Files

Supplemental File 1 - Annotations

This TSV file contains all annotations including superclass, cell class, cell type, hemilineage, side, neurotransmitter and representative coordinates.

Supplemental File 2 - Summary with NGL links

This CSV file contains a summary of all hemilineages including neuroglancer links for viewing.

Supplemental File 3 - Hemilineage clustering

This CSV file contains details on the clustering of hemilineages into morphology groups.

Supplemental File 4 - Hemibrain metadata

This CSV file contains metadata for hemibrain neurons including columns for soma *side* and *cell class*.

## Supplemental Videos

Supplemental Video 1

3D rendering showing all FlyWire neurons.

Supplemental Video 2

3D rendering showing all FlyWire neurons by superclass.

Supplemental Video 3

Slideshow for morphology groups by hemi-lineage.

Supplemental Video 4

Slideshow of hemibrain-FlyWire matched cell types.

## Methods

### Data and Tool availability

A static dump of data artefacts from this paper is available at https://github.com/flyconnectome/flywire_annotations. This includes:

- neuron annotations + other metadata
- high quality skeletons for all proofread FlyWire neurons
- NBLAST scores for FlyWire vs. hemibrain
- NBLAST scores for FlyWire left vs. right

Annotations are additionally available via Codex (https://codex.flywire.ai/), the connectome annotation versioning engine (CAVE) which can be queried through e.g. the CAVEclient (https://github.com/seung-lab/CAVEclient), and the FAFB-FlyWire CATMAID spaces (https://fafb-flywire.catmaid.org). At the time of writing access to Codex and CAVE require signing up using a Google account.

To aid a number of analyses, hemibrain meshes were mapped into FlyWire (FAFB14.1) space. These can be co-visualised within neuroglancer for example by following this link: https://tinyurl.com/flywirehb. This also enables direct querying of both our flywire annotations and hemibrain annotations from within neuroglancer to efficiently find and compare cell types. This scene also includes a second copy of the hemibrain data (layer `hemibrain_meshes_mirr`) which has been non-rigidly mapped onto the opposite side of FAFB.

Analyses were performed using open source packages using both the R natverse^104^ and Python navis infrastructures (see Table 1). The fafbseg R and Python packages have extensive functionality dedicated to working with FlyWire data, including fetching connectivity and querying the segmentation. Unless otherwise stated, all analyses were performed against the 630 release version (i.e. the first public data release for Flywire).

**Table 1:**
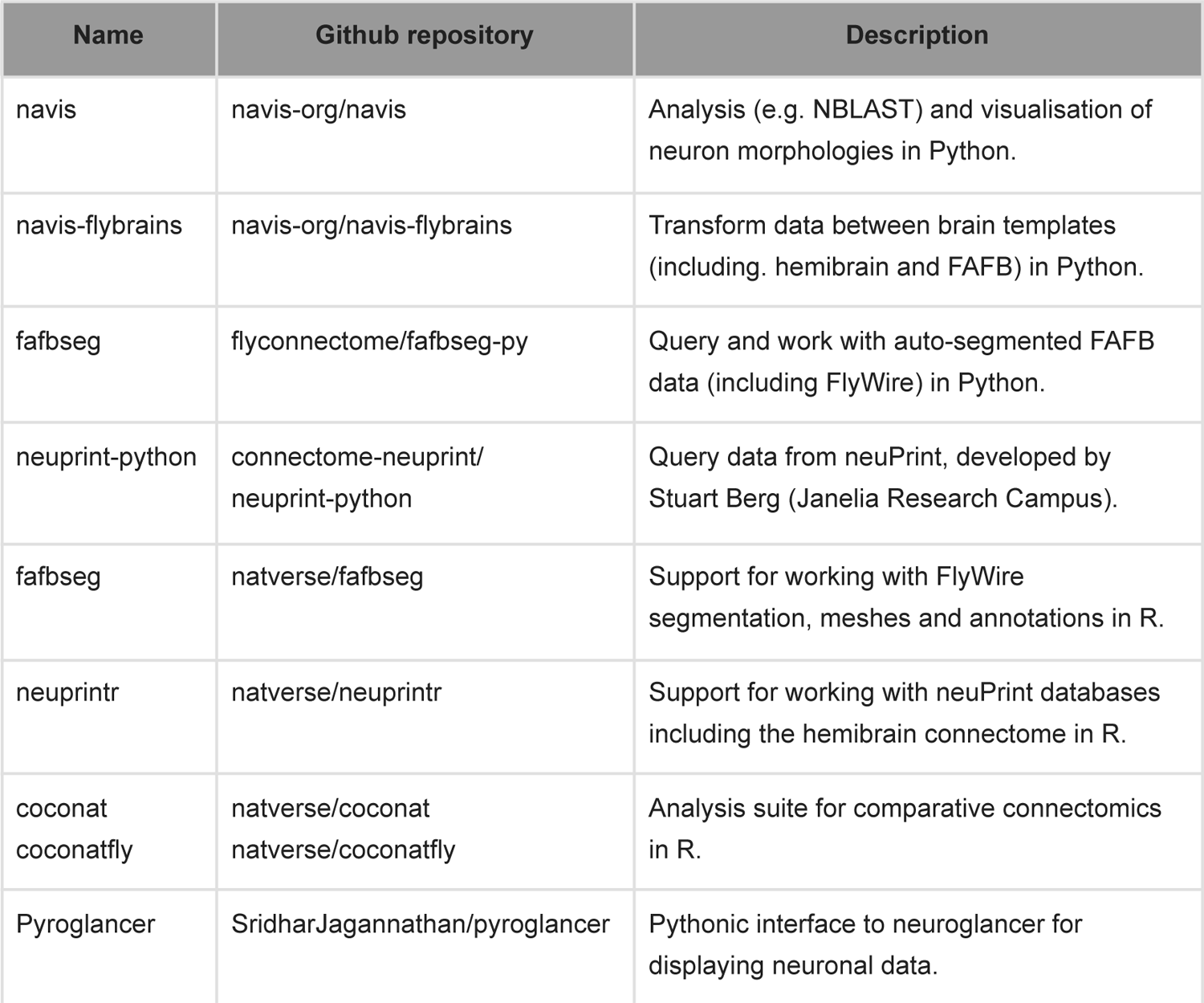
Software tools used.

### Annotations

#### Base annotations

At the time of writing, the general FlyWire annotation system operates in a read-only mode in which users can add additional annotations for a neuron but cannot edit or delete existing annotations. Furthermore, the annotations consist of a single free-form text field bound to a spatial location. This enabled many flywire users (including our own group) to contribute a wide range of *community* annotations, which are reported in our companion paper^1^ but are not considered in this study. As it became apparent that a complete connectome could be obtained, we found that this approach was not a good fit for our goal of obtaining a structured, systematic and canonical set of annotations for each neuron with extensive manual curation. We therefore set up a web database (seatable, see https://seatable.io/) that allowed records for each neuron to be edited and corrected over time; columns with specific acceptable values were added as necessary.

Each neuron was defined by a single point location (aka root point) and its associated PyChunkedGraph supervoxel. Root IDs were updated every 30min by a Python script based on the fafbseg package (see Table 1) to account for any edits. The canonical point for the neuron was either a location on a large calibre neurite within the main arbour of the neuron, a location on the cell body fibre close to where it entered the neuropil, or a position within the nucleus as defined by the nucleus segmentation table^105^. The former was preferred since segmentation errors in the cell body fibre tracts regularly resulted in the wrong soma being attached to a given neuronal arbour. These soma swap errors persisted late into proofreading and when fixed resulted in annotation information being attached to the wrong neuron until this in turn was fixed.

We also note that our annotations include a number of non-neuronal cells/objects such as glia cells, trachea, ECM which others might find useful (superclass *not_a_neuron*).

#### Soma Position and Side

Besides the canonical root point, the soma position was recorded for all neurons with a cell body. This was either based on curating entries in the nucleus segmentation table (removing duplicates or positions outside the nucleus) or on selecting a location, especially when the cell body fibre was truncated and no soma could be identified in the dataset. These soma locations were critical for a number of analyses and also allowed a consistent *side* to be defined for each neuron. This was initialised by mapping all soma positions to the symmetric JRC2018F template and then using a cutting plane at the midline perpendicular to the mediolateral (X) axis to define left and right. However all soma positions within 20 µm of the midline plane were then manually reviewed. The goal was to define a consistent logical soma side based on examination of the cell body fibre tracts entering the brain; this ultimately ensured that cell types present e.g. in one copy per brain hemisphere were always annotated so that one neuron was identified as the left and the other the right. In a small number of cases - e.g. for the bilaterally symmetric octopaminergic VUM (ventral unpaired medial) neurons - we assigned *side* “central”.

For sensory neurons, *side* refers to whether they enter the brain through the left or the right nerve. In a small number of cases we could not unambiguously identify the nerve entry side and assigned *side* “na”.

#### Hierarchical annotations

Hierarchical annotations include *flow*, *superclass*, *class* (+ a *subclass* in certain cases) and *cell type*. The *flow* and *superclass* was generally assigned based on an initial semi-automated approach followed by extensive and iterative manual curation. See Supplemental Table S2 for definitions and sections below for details on certain superclasses.

Based on the superclasses we define two useful groupings which are used throughout the main text:

- *Central brain neurons* consist of all neurons with their somata in the central brain defined by the five superclasses: *central, descending, visual centrifugal, motor, endocrine*.
- *Central brain associated neurons* further include superclasses: *visual projection* neurons, *ascending* neurons and *sensory* neurons (but omit sensory neurons with cell class: *visual*).

Cell *classes* represent salient groupings/terms that have been previously used in the literature (see Supplemental Table S2 for examples). For sensory neurons, the *class* indicates their modality (where known).

#### Sensory and motor neuron identification

We identified all non-visual sensory and motor neurons entering/exiting the brain through the antennal, eye, occipital, and labial nerves by screening all axon profiles in a given nerve.

Sensory neurons were further cross-referenced to existing literature to assign modalities (via the *class* field) and, where applicable, a *cell type*. Previous studies have identified almost all head mechanosensory bristle and taste peg mechanosensory neurons (BMN and TPMN; ^106^) in the left hemisphere (at time of publication: right hemisphere). Gustatory sensory neurons (GRN) were previously identified by ^107^ and Johnston’s Organ neurons (JON) by ^108, 109^ and ^109^ in a version of FAFB that employed manual reconstruction (https://fafb.catmaid.virtualflybrain.org). Those neurons were identified in the FlyWire instance by transformation and overlay onto FlyWire space as described by ^106^.

JONs in the right hemisphere were characterised based on innervation of the major AMMC zones (A, B, C, D, E and F), but not further classified into subzone innervation as shown in^108^. Other sensory neurons (BMNs, TPMNs and GRNs) in the right hemisphere were identified through NBLAST-based matching of their mirrored morphology to the left hemisphere and expert review. Olfactory, thermo- and hygrosensory of the antennal lobes were identified via their connectivity to cognate uniglomerular projection neurons and NBLAST-based matching to previously identified hemibrain neurons^60, 110^.

Visual sensory neurons (R1-6, R7-8 and ocellar photoreceptor neurons) were identified by manually screening neurons with presynapse in either the lamina, the medulla and/or the ocellar ganglia^111^. Out of the expected 12,800 photoreceptor neurons from the retina we have found only 3,943 so far due but more will be identified in the future. For the ocelli, all 270 photoreceptor neurons were annotated.

#### Ascending and descending neuron identification

We seeded all profiles in a cross section in the ventral posterior gnathal ganglion (GNG) through the cervical connective to identify all neurons entering and exiting the brain at the neck. We identified all descending neurons (DNs) based on the following criteria:

1. Soma located within the brain dataset
2. Main axon branch leaving the brain through the cervical connective

We then classified the DNs based on their soma location according to ^112^. Briefly, the soma of DNa, DNb, DNc and DNd is located in the anterior half (a - anterior dorsal, b - anterior ventral, c - in the pars intercerebralis, d - outside cell cluster on the surface) and DNp in the posterior half of the central brain. DNg somas are located in the GNG.

To identify DNs described by ^112^ in the EM dataset we transformed the volume renderings of DN GAL4 lines into FAFB space. Displaying EM and LM neurons in the same space enabled accurate matching of closely morphologically related neurons. For DNs without available volume renderings we identified candidate EM matches by eye, transformed them into JRC2018U space and overlaid them onto the GAL4 or Split GAL4 line stacks (named in ^112^ for that type) in FIJI for verification. With these methods we identified all but 2 (DNd01 and DNg25) in FAFB and annotated their cell type with the published nomenclature. All other unmatched DNs received a systematic cell type consisting of their soma location, an “e” for EM type and a three digit number (e.g. DNae001). A detailed account and analysis of descending neurons will be published separately (Eichler *et al.*, in prep).

#### Hemilineage annotations

Central nervous system lineages were initially mapped for the 3rd instar larval brain, where, for each lineage, the neuroblast of origin and its progeny are directly visible^113–116^. Genetic tools that allow stochastic clonal analysis^117^ have enabled researchers to visualise individual lineages as GFP-marked “clones”. Clones reveal the stereotyped morphological footprint of a lineage, its overall “projection envelope”^47^, as well as the cohesive fibre bundles - so-called “hemilineage-associated tracts” (HATs) - formed by neurons belonging to it. Using these characteristics, lineages could be also identified in the embryo and early larva^118, 119^, as well as in pupae and adults^46–49, 53, 120^. HATs can be readily identified in the EM image data, and we used them, in conjunction with clonal projection envelopes, to identify hemilineages in the EM dataset through a combination of the following methods:

1. Visual comparison of HATs formed by reconstructed neurons in the EM, and the light microscopic map reconstructed from anti-Neuroglian-labelled brains^46, 48, 49^: In cross section, tracts typically appear as clusters of 50-100 tightly packed, rounded contours of uniform diameter (∼200 nm), surrounded by neuronal cell bodies (when sectioned in the cortex) or irregularly shaped terminal neurite branches and synapses (when sectioned in the neuropil area; Figure 2C). The point of entry and trajectory of a HAT in the neuropil is characteristic for a hemilineage.
2. Matching branching pattern of reconstructed neurons with the projection envelope of clones: as expected from the light microscopic map based on anti-Neuroglian-labelled brains^46^, the majority of hemilineage tracts visible in the EM dataset occur in pairs or small groups (3-5). Within these groups, individual tracts are often lined by fibres of larger (and more variable) diameter, as shown in Figure 2C. However, the boundary between closely adjacent hemilineage tracts is often difficult to draw based on the EM image alone. In these cases, visual inspection and quantitative comparison of the reconstructed neurons belonging to a hemilineage tract with the projection envelope of the corresponding clone, which can be projected into the EM dataset through Pyroglancer (see Table 1), assists in properly assigning neurons to their hemilineages.
3. Identifying homologous HATs across three different hemispheres (left and right of FlyWire, hemibrain): by comparison of morphology (NBLAST^54^), as well as connectivity (assuming that homologous neurons share synaptic partners), we were able to assign the large majority of neurons to specific HATs that matched in all three hemispheres.

In existing literature two systems for hemilineage nomenclature are used: “Ito/Lee”^48, 49^ and “Hartenstein”^46, 47^. While they overlap in large parts, some lineages have only been described in one but not the other nomenclature. In the main text, we provide (hemi-)lineages according to the “ItoLee” nomenclature for simplicity. Below and in the supplements we also provide both names as “ItoLee/Hartenstein”, and the mapping between the two nomenclatures is provided in <u>Supplementary File 2</u>. From prior literature we expected a total of around 120 lineages in the central brain, including the gnathal ganglia (GNG)^46–49, 103^. Indeed, we were able to identify 120 lineages based on light-level clones and tracts, as well as the HATs in FlyWire. Among these, LHp3/CP5 could not be matched to any clone. We have not yet identified SLPpm4 which is only present in^48^. Therefore, together, we expect to eventually identify 121 lineages.

By comprehensively inspecting the hemilineage tracts originally in CATMAID and then in FlyWire, we can now reconcile previous reports. Specifically, new to^48, 49^ (‘ItoLee nomenclature’) are: CREl1/DALv3, LHp3/CP5, DILP/DILP, LALa1/BAlp2, SMPpm1/DPMm2 and VLPl5/BLVa3_or_4 - we gave these neurons lineage names following the naming scheme in^48, 49^. New to^46^ (‘Hartenstein nomenclature’) are: SLPal5/BLAd5, SLPav3/BLVa2a, LHl3/BLVa2b, SLPpl3/BLVa2c, PBp1/CM6, SLPpl2/CP6, SMPpd2/DPLc6, PSp1/DPMl2, and LHp3/CP5 - we named these units following the Hartenstein nomenclature naming scheme. We did not take the following clones from^48^ into account for the total count of lineages/hemilineages, because they originate in the optic lobe and their neuroblast of origin has not been clearly demonstrated in the larva: VPNd2, VPNd3, VPNd4, VPNp2, VPNp3, VPNp4, VPNv1, VPNv2, VPNv3.

For calculating the total number of hemilineages, to keep the inclusion criteria consistent with the lineages, we included the Type II lineages (DL1-2/CP2-3, DM1-6 / DPMm1, DPMpm1, DPMpm2, CM4, CM1, CM3) by counting the number of cell body fibre tracts, acknowledging that they may or may not be hemilineages. Neuroblasts of type II lineages, instead of generating ganglion mother cells that each divide once, amplify their number, generating multiple intermediate progenitors (IPs) that in turn continue dividing like neuroblasts^43, 121, 122^. It has not been established how the tracts visible in type II clones (and included in <u>Figure S2.1</u> and <u>Supplemental Files 2 and 3</u>) relate to the (large number of) type II hemilineages.

There are also 3 type I lineages (VPNl&d1/BLAl2, VLPl2/BLAv2 and VLPp&l1/DPLpv) with more than two tracts in the clone; we included these additional tracts in the hemilineages provided in the text. Without taking these Type I and Type II tracts into account, we identified 141 hemilineages.

A minority of neurons in the central brain could not reliably be assigned to a lineage. These mainly include the (putative) primary neurons (3,614). Primary neurons, born in the embryo and already differentiated in the larva, form small tracts with which the secondary neurons become closely associated^52^. In the adult brain, morphological criteria that unambiguously differentiate between primary and secondary neurons have not yet been established. In cases where experimental evidence exists^42^ primary neurons have significantly larger cell bodies and cell body fibres. Loosely taking these criteria into account we surmise that a fraction of primary neurons form part of the HATs defined as described above. However, aside from the HATs, we see multiple small bundles, typically close to but not contiguous with the HATs, which we assume to consist of primary neurons. Overall, these small bundles contained 3,614 neurons, designated as “primary” or “putative primary” neurons.

#### Morphological groups

We further subdivided hemilineages into groups of neurons defined by shared morphological characteristics (“morphological groups”). As a first step, neurons were mirrored from one hemisphere to another, and pairwise morphological similarity scores were generated using NBLAST^54^. Neurons were then hierarchically clustered. To determine the coarseness of the groups (i.e. where to cut the dendrogram exemplified in Figure 2I), we used the ‘elbow method’^123^, which uses the point of the most abrupt change in the cluster distance. Morphological groups were named after the compartments that they innervated (Supplemental files 2-3). Figure 2I illustrates the classification of hemilineage AOTUv3_dorsal, for which our method defines four groups:

1. The first group (yellow in Fig 2I, AOTUv3_dorsal_1/AOTU_BU_MB_PED in <u>Suppl.</u> <u>File 3</u>) innervates a small domain in the lateral AOTU and projects to the dorsal and anterior bulb (BU; yellow); this class corresponds to the previously defined TuBu_s and TuBu_a neurons^124, 125^.
2. The second group (green in Fig 2I, AOTUv3_dorsal_4/AOTU_LAL_SMP in <u>Suppl.</u> <u>File 3</u>) also has branches in medial AOTU and SIP SMP, but projects to the LAL.
3. The third (red in Fig 2I, AOTUv3_dorsal_3/AOTU_IB_SIP in <u>Suppl. File 3</u>) has proximal branches in the medial AOTU and (sparsely) the SIP, and projects to the inferior bridge (IB), as well as to the contralateral AOTU and SIP.
4. The fourth (purple in Fig 2I, AOTUv3_dorsal_2/IB_SIP_SMP in <u>Suppl. File 3</u>) lacks innervation of the AOTU, and interconnects the adjacent SIP with SMP and IB.

In a similar manner, other hemilineages cluster into discrete non-overlapping groups of neurons that represent anatomical “modules”, forming connections between specific brain domains. The morphological groups, and the top three neuropils innervated by each morphological group, are recorded in <u>Supplementary Files 2 and 3</u> (at the time of writing the groups are not yet visible in codex.flywire.ai). There are in general 2-5 (10/90 percentile) morphological groups per hemilineage; and 3-44 neurons per morphological group.

### FAFB Laterality

Our observations (and those of the Rachel Wilson lab, Pratyush Kandimalla and Stephane Noselli, see Acknowledgements) of the asymmetric body (AB) in FAFB-FlyWire led us to catch an error made in assembling the original FAFB14 dataset^32^. The asymmetric bodies of the central complex should be larger on the fly’s right^56, 57^ but in the FAFB volume appeared enlarged on the left. Although we initially considered the possibility of situs inversus, this appears to be extremely rare^58, 126^. By going back to the original grids that had been prepared for EM and comparing ultrastructural features to the registered image data set we conclude instead that FAFB14 was digitally inverted during image acquisition. Unfortunately at this point in time mirroring the FAFB14 volume, FlyWire segmentation and other associated data would introduce many complications for analysis. We therefore corrected the ‘left’ and ‘right’ labels so that they match the true biological side; these are used throughout this work, all other FlyWire papers and used in codex.flywire.ai and the FAFB-FlyWire CATMAID space. For consistency with the data set as it can be viewed in neuroglancer or other viewers and with previously published FAFB work, images in this paper and its companion papers retain the original image orientation i.e. they are mirrored relative to the ‘true’ fly brain (Figure 2K; <u>Figure S2.2</u>).

For neuronal data that might be asymmetric, the dataset can still be mirrored for display or analysis e.g. when comparing with other data sets, such as the hemibrain. We have built tools to enable users to digitally mirror FAFB-FlyWire data using the Python <u>flybrains</u> or natverse <u>nat.jrcbrains</u> packages (<u>Figure S2.2C</u>). In more detail, since the hemibrain and FAFB datasets are different shapes, a non-rigid transform already exists between them. This passes via the symmetric JRC2018F light level template^127^. Therefore to superimpose the (true) RHS of hemibrain and FAFB you can use the existing transforms, but apply a simple mirror flip within the JRC2018F space. The `nat.jrcbrains::mirror_fafb()` function does FAFB→JRC2018F→horizontal flip in JRC1018F→FAFB. The `navis.mirror_brain` function applies almost exactly the same transformation by fitting a thin plate spline landmarks transformation.

We also provide a neuroglancer scene in which both flywire and hemibrain data is displayed in the correct orientation: https://tinyurl.com/flywirehbflip. The hemibrain neurons have been non-rigidly mapped onto FAFB’s (true) RHS and then both FAFB and hemibrain data have been flipped along the X axis. In this scene a frontal view has both FAFB and hemibrain (true) RHS to the left of the screen. By default the scene displays the SA1 and SA2 neurons which target the right asymmetric body for both flywire and the hemibrain, confirming that the RHS for both datasets has been superimposed. Note that any coordinates recorded in this scene are necessarily different from coordinates based on the standard orientation of FAFB.

### Morphological comparisons

Throughout our analyses we make use of NBLAST^54^ to generate morphological similarity scores between neurons - e.g. for matching neurons between the FlyWire and the hemibrain datasets, or for the morphological clustering of the hemilineages. In brief, NBLAST treats neurons as point clouds with associated tangent vectors describing directionality, so called “dotprops”. For a given query→target neuron pair, we perform a KNN search between the two point clouds and score each nearest-neighbour pair by their distance and the dot product of their vector. These are then summed up to compute the final query→target NBLAST score. It is important to note that direction of the NBLAST matters, i.e. NBLASTing neurons A→B ≠ B→A. Unless otherwise noted, we use the mean of the forward and reverse NBLAST scores.

The NBLAST algorithm is implemented in both *navis* and the *natverse* (see Table 1). However, we modified the *navis* implementation for more efficient parallel computation in order to scale to pools of more than 100k neurons. For example, the all-by-all NBLAST matrix for the full 128k Flywire neurons alone occupies over 500GB of memory (32bit floats). Most of the large NBLASTs were run on a single cluster node with 112 CPUs and 1TB RAM provided by the MRC LMB Scientific Computing group, and took between 1 and 2 days (wall time) to complete.

Below we provide recipes for the different NBLAST analyses used in this paper:

### FlyWire all-by-all NBLAST

For this NBLAST, we first generated skeletons using the L2 cache. In brief: underlying the Flywire segmentation is an octree data structure where level 0 represents supervoxels which are then agglomerated over higher levels^128^. The second layer (L2) in this octree represents neurons as chunks of roughly 4×4×10μm in size which is sufficiently detailed for NBLAST. The L2 cache holds precomputed information for each L2 chunk, including a representative x/y/z coordinate in space. We used the x/y/z coordinates and connectivity between chunks to generate skeletons for all FlyWire neurons (implemented in *fafbseg*; see Table 1). Skeletons were then pruned to remove side branches smaller than 5 microns. From those skeletons we generated the dotprops for NBLAST using *navis*.

Before the NBLAST, we additionally transformed dotprops to the same space by mirroring those from neurons with *side* right onto the left. The NBLAST was then run only in forward direction (query→target) but because the resulting matrix was symmetrical we could generate mean NBLAST scores using the transposed matrix: (A + A^T^) / 2. This NBLAST was used to find left/right neuron pairs, define (hemi-)lineages and run the morphology group clustering.

### FlyWire - hemibrain NBLAST

For FlyWire, we re-used the dotprops generated for the all-by-all NBLAST (see previous section). To account for the truncation of neurons in the hemibrain volume, we removed points that fell outside the hemibrain bounding box.

For the hemibrain, we downloaded skeletons for all neurons from neuPrint (https://neuprint.janelia.org) using *neuprint-python* and *navis* (see Table 1). In addition to the ∼23k typed neurons we also included all untyped neurons (often just fragments) for a total of 98k skeletons. These skeletons were pruned to remove twigs smaller than 5 microns and then transformed from hemibrain into FlyWire (FAFB14.1) space using a combination of non-rigid transforms^127, 128^ (implemented via *navis*, *navis-flybrain* and *fafbseg*; see Table 1). Once in FlyWire space, they were resampled to 0.5 nodes per micron of cable to approximately match the resolution of the FlyWire L2 skeletons, and then turned into dotprops. The NBLAST was then run both in forward (FlyWire→hemibrain) and reverse (hemibrain→FlyWire) direction and the mean scores were used.

This NBLAST allowed us to match FlyWire left against the hemibrain neurons. To also allow matching FlyWire right against the hemibrain, we performed a second run after mirroring the FlyWire dotprops to the opposite side.

### Cross-brain co-clustering

The pipeline for the morphology-based across brain co-clustering used in Figure 6 was essentially the same as for the FlyWire - hemibrain NBLAST with two exceptions:

1. We used high-resolution FlyWire skeletons instead of the coarser L2 skeletons (see below).
2. Both FlyWire and hemibrain skeletons were resampled to 1 node per micron before generating dotprops.

### High resolution skeletonisation

In addition to the coarse L2 skeletons, we also generated high-res skeletons which were, for example, used to calculate the total length of neuronal cable reported in our companion paper^1^ (146m). Briefly, we downloaded neuron meshes (LOD 1) from the flat 630 segmentation (available at gs://flywire_v141_m630) and skeletonised them using the “wavefront” method implemented in *skeletor* (https://github.com/navis-org/skeletor). Skeletons were then rerooted to their soma (if applicable), smoothed (by removing small artifactual bristles on the backbone), healed (segmentation issues can cause breaks in the meshes) and slightly downsampled. A modified version of this pipeline is implemented in *fafbseg*. Skeletons are available for download (see section on Data and Tool availability).

### Connectivity normalisation

Throughout this paper, the basic measure of connection strength is the number of unitary synapses between two or more neurons^129^; connections between adult fly neurons can reach thousands of such unitary synapses^3^. Previous work in larval *Drosophila* has indicated that synaptic counts approximate contact area^130^ which is most commonly used in mammalian species when a high resolution measure of anatomical connection strength is required. Connectomics studies also routinely use connection strength normalised to the target cell’s total inputs^91, 129^. For example, if neurons *i* and *j* are connected by 10 synapses and neuron *j* receives 200 inputs in total, the normalised connection weight *i→j* would be 5%. Gerhard et al. (2017) showed that while absolute number of synapses for a given connection changes drastically over the course of larval stages, the proportional (i.e. normalised) input to the downstream neuron remains relatively constant^101^. Importantly, we have some evidence (Figure 4F) that normalised connection weights are robust against technical noise (differences in reconstruction status, synapse detection). Note that for analyses of MB circuits we use an approach based on the fraction of the input or output synaptic budget associated with different KC cell types; this differs slightly from the above definition and will be detailed in a separate section below.

### Cell typing

Most of our cell type annotations (72%) are derived from the hemibrain cell type matching effort (see section below). The remainder was generated either by comparing to existing literature (e.g. in case of some optic lobe cell types or sensory neurons) and/or by finding unambiguous left/right pairs through a combination of NBLAST and connectivity clustering (<u>Figure S8</u>). New types without pre-existing name were given a simple numerical cross-brain identifier (e.g. “*CB0001*”) as a (temporary) cell type label.

For provenance, we provide two columns of cell types in our supplemental data:

- *hemibrain_type* always refers to one or more hemibrain cell types; in rare occasions where a matched hemibrain neuron did not have a type we recorded body IDs instead
- *cell_type* contains types that are either not derived from the hemibrain or that represent refinements (e.g. a split or retyping) of hemibrain types Neurons can have both a *cell_type* and a *hemibrain_type* entry in which case the *cell_type* represents a refinement or correction and should take precedence.

### Hemibrain cell type matching

We used NBLAST^54^ to match FlyWire neurons to hemibrain cell types (see section on <u>NBLASTs</u> for details). From the NBLAST scores we extracted, for each FlyWire neuron, a list of potential cell type hits using all hits in the 90th percentile. Individual FlyWire neurons were co-visualized with their potential hits in neuroglancer (see link in <u>Data and Tool availability</u> <u>section</u>) and the correct hit (if found) was recorded. In difficult cases, we would also inspect the subtree of the NBLAST dendrograms containing the neurons in questions to include local cluster structure in the decision making (<u>Figure S3</u>). In cases where two or more hemibrain cell types could not be cleanly delineated in FlyWire (i.e. there were no corresponding separable clusters) we recorded “composite” (many:1) type matches (Figure 3E; <u>Figure</u> <u>S3G</u>; <u>Figures S8</u>).

When a matched type was either missing large parts of its arbours due to truncation in the hemibrain or the comparison with the FlyWire matches suggested closer inspection was required, we used cross-brain connectivity comparisons (see <u>section below</u>) to decide whether to adjust (split or merge) the type. A merge of two or more hemibrain types was recorded as e.g. “SIP078,SIP080” while a split would be recorded as “PS090a” and “PS090b” (i.e. with a lower-case letter as suffix). In rare cases where we were able to find a match for an untyped hemibrain neuron, we would record the hemibrain body ID as hemibrain type and assign a CBXXXX identifier as cell type.

Finally, the hemibrain introduced the concept of “morphology types” and “connectivity types”^3^. The latter represent refinements of the former and differ only in their connectivity. For example, morphology type *SAD051* splits into two connectivity types: *SAD051_a* and *SAD051_b*, where the *_{letter}* indicates that these are connectivity types. Throughout our FlyWire↔hemibrain matching efforts we found connectivity types hard to reproduce and our default approach was to match only up to the morphology type. In some cases, e.g. antennal lobe local neuron types like *lLN2P_a* and *lLN2P_b,* we were able to find the corresponding neurons in FlyWire.

We note that in numerous cases that we reviewed but remain unmatched we encountered what we call ambiguous “daisy-chains”: imagine 4 fairly similar cell types, *A*, *B*, *C*, and *D*. Often these adjacent cell types represent a spectrum of morphologies where *A* is similar to *B*, *B* is similar to *C* and *C* is similar to *D*. The problem now is in unambiguously telling A from B, B from C and C from D. But at the same time *A* and *D* (on opposite ends of the spectrum) are so dissimilar that we would not expect to assign them the same cell type. These kinds of graded or continuous variation have been observed in a number of locations in the mammalian nervous system and represent one of the classic complications of cell typing^33^. Absent other compelling information that can clearly separate these groups, the only reasonable option would seem to be to lump them together. Since this would erase numerous proposed hemibrain cell types, the de facto standard for the fly brain, we have been conservative about making these changes pending analysis of additional connectome data^3^.

### Hemibrain cell type matching with connectivity

In our hemibrain type matching efforts about 12% of cell types could not be matched 1:1. In these cases we used across-dataset connectivity clustering (e.g. to confirm the split of a hemibrain type or a merger of multiple cell types). To generate distances, we first produced separate adjacency matrices for each of the three hemispheres (FlyWire left, right and hemibrain). In these matrices, each row is a query neuron and each column is an up- or downstream *cell type*; the values are the connection weights (i.e. number of synapses). We then combine the three matrices along the first axis (rows) and keep only the cell types (columns) which have been cross-identified in all hemispheres. From the resulting observation vector we calculate a pairwise cosine distance. It is important to note that this connectivity clustering depends absolutely on the existence of a corpus of shared labels between the two datasets – without such shared labels, which were initially defined by morphological matching as described above, connectivity matching cannot function.

This pipeline is implemented in the *coconatfly* package (see Table 1), which provides a streamlined interface to carry out such clustering. For example the following command can be used to see if the types given to a selection of neurons in the Lateral Accessory Lobe (LAL) are robust.

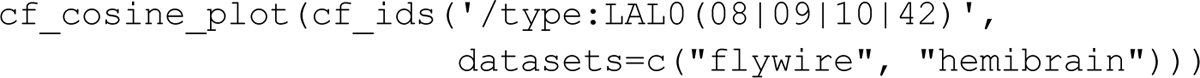

An optional interactive mode allows for efficient exploration within a web browser. For further details and examples, see https://natverse.org/coconatfly/.

### Defining robust cross-brain cell types

In Figure 6 we used two kinds of distance metrics: one calculated from connectivity alone (used for FC1-3; Figure 6E-G) and a second combining morphology + connectivity (used for FB1-9; Figure 6H-K) to help define robust cross-brain cell types. The connectivity distance is as described <u>above</u>. The combined morphology + connectivity distances were generated by taking the sum of the connectivity and NBLAST distances. Connectivity-only works well in case of cell types that don’t overlap in space but instead tile a neuropil. For cell types that are expected to overlap in space we find that adding NBLAST distances is a useful constraint to avoid mixing of otherwise clearly different cells. From the distances we generated a dendrogram representation using the Ward algorithm and then extracted the smallest possible clusters that satisfy two criteria:

The resulting groups were then manually reviewed (Figure S8).

### Availability via CATMAID Spaces

To increase the accessibility and reach of the annotated FlyWire connectome, meshes of proofread FlyWire neurons and synapses were skeletonized and imported into CATMAID, a widely used web-based tool for collaborative tracing, annotation, and analysis of large-scale neuronal anatomy datasets (https://catmaid.org129,131; <u>Figure S7</u>). Spatial annotations like skeletons are modelled using PostGIS data types, a PostgreSQL extension that is popular in the Geographic Information System community. This allows us to reuse many existing tools to work with large spatial datasets, e.g. indexes, spatial queries and mesh representation.

A publicly available version of the FlyWire CATMAID project is available at https://fafb-flywire.catmaid.org. This project leverages a new extension, called CATMAID Spaces (https://catmaid.org/en/latest/spaces.html), which allows users to create and administer their own tracing and annotation environments on top of publicly available neuronal image volumes and connectomic datasets. In addition, users can now login through the public authentication service ORCiD (https://www.orcid.org), so that everyone can log-in on public CATMAID projects. Users can also now create personal copies (“Spaces”) of public projects. The user then becomes an administrator, and can invite other users, along with the management of their permissions in this new project. Invitations are managed through project tokens, which the administrator can generate and send to invitees for access to the project. Additional publicly available resources for use in CATMAID Spaces can be found at https://spaces.catmaid.org. Both CATMAID platforms can talk to each other and it is possible to load data from the dedicated FAFB-FlyWire server in the more general Spaces environment.

Metadata annotations for each neuron (root id, cell type, hemilineage, neurotransmitter) were imported for FlyWire project release 630. Skeletons for all 127,978 proofread neurons were generated from the volumetric meshes (see <u>Methods section on skeletonization</u>), downsampled (ds=4), and imported into CATMAID, resulting in 177,418,713 treenodes. To reduce import time, skeletons were imported into CATMAID directly as database inserts via SQL, rather than through public RESTful APIs. FlyWire root IDs are available as metadata for each neuron, facilitating interchange with related resources such as FlyWire Codex^1^. Synapses attached to reconstructed neurons were imported as CATMAID connector objects and attached to neuron skeletons by doing a PostgreSQL query to find the nearest node on each of the partner skeletons. Connector objects only were linked to postsynaptic partners if the downstream neuron was in the proofread data release (177,471,255 connections from the 130,105,549 synapses with at least one partner in the proofread set).

### Synapse counts

Insect synapses are polyadic, i.e. each presynaptic site can be associated with multiple postsynaptic sites. In contrast to the Janelia hemibrain dataset, the synapse predictions used in FlyWire do not have a concept of a unitary presynaptic site associated with a T-bar^64^. Therefore, presynapse counts used in this paper do not represent the number of presynaptic sites but rather the number of outgoing connections.

In *Drosophila* connectomes, reported counts of the inputs (postsynapses) onto a given neuron are typically lower than the true number. This is because fine-calibre dendritic fragments frequently cannot be joined onto the rest of the neuron, instead remaining as free-floating fragments in the dataset.

### Technical noise model

To model the impact of technical noise such as proofreading status and synapse detection on connectivity, we first generated a fictive “100%” ground-truth connectivity. We took the connectivity between cell-typed left FlyWire neurons and scaled each edge weight (=number of synapses) by the postsynaptic completion rates in the respective neuropil. For example, all edge weights in the left mushroom body calyx (CA) which has a postsynaptic completion rate of 45.8% were scaled by a factor of 100/45.8=2.18.

In the second step, we simulated the proofreading process by randomly drawing (without replacement) individual synaptic connections from the fictive ground-truth until reaching a target completion rate. We further simulate the impact of false-positive and false-negatives by randomly adding and removing synapses to/from the draw according to the precision (0.72) and recall (0.77) rates reported by Buhmann *et al.*^64^. In each round, we made two draws:

1. A draw using the original per-neuropil postsynaptic completion rates.
2. A draw where we flip the completion rates for left and right neuropils, i.e. use the left CA completion rate for the right CA and *vice versa*.

In each of the 500 rounds we ran, we drew two weights for each edge. Both stem from the same 100% ground-truth connectivity but have been drawn according to the differences in left vs right hemisphere completion rates. Combining these values we calculated the mean difference and quantiles as function of the weight for the Flywire left (i.e. the draw that was not flipped) (Figure 5G). We focussed this analysis on edge weights between 1 and 30 synapses because the number of edges stronger than that is comparatively low, leaving gaps in the data.

### Kenyon cell analyses

#### Connection weight normalisation and synaptic budget analysis

When normalising connection weights, we typically convert them to the percentage of total input onto a given target cell (or cell type). However, in the case of the mushroom body the situation is complicated by what we think is a technical bias in the synapse detection methods used for the two connectomes which causes certain kinds of unusual connections to be very different in frequency between the two datasets. We find that the total number of postsynapses as well as the postsynapse density in the mushroom body lobes are more than doubled in the hemibrain compared to Flywire (<u>Figure S5B,C</u>). This appears to be explained by certain connections (especially KC→KC connections, which are predominantly arranged with an unusual “rosette” configuration along axons and whose functional significance is poorly understood^132^) being much more prevalent in the hemibrain than in FlyWire (<u>Figure S5D</u>). Some other neurons including the APL giant interneuron also make about twice as many synapses onto Kenyon Cells (KC) in the hemibrain compared with FlyWire (<u>Figure S5A</u>). As a consequence of this large number of inputs onto KC axons in the hemibrain, input percentages from all other cells are reduced in comparison with FlyWire.

To avoid this bias, and because our main goal in the KC analysis was to compare different populations of Kenyon cells, we instead expressed connectivity as a fraction of the total synaptic budget for upstream or downstream cell types. For example, we asked, “what fraction of the APL’s output is spent on each of the different KC types”? Similarly, we quantified connectivity for individual KCs as a fraction of the budget for the whole KC population.

### Calculating *K* from observed connectivity

Calculation of *K*, i.e. the number of unique odour channels that each Kenyon Cell receives input from, was principally based on their synaptic connectivity. For this, we looked at their inputs from uniglomerular antennal lobe projection neurons (ALPNs) and asked from how many of the 58 antennal lobe glomeruli does a KC receive input from. *K* as reported in Figure 6 is based on non-thresholded connectivity. Filtering out weak connections does lower *K* but importantly, our observations (e.g. that KCg-m have a lower *K* in FlyWire than in the hemibrain) are stable across thresholds (<u>Figure S5G</u>).

### Kenyon cell model

A simple rate model of neural networks (Litwin-Kumar et al. 2017) was used to generate the theoretical predictions of K, the number of ALPN inputs that each KC receives (seen in Figure 5K). KC activity is modelled by

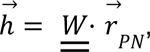

where ℎ is a vector of length *M* representing KC activity, *W* is an *M* × *N* matrix representing the synaptic weights between the KCs and PNs, *r* is a vector of length *N* representing PN activity. The number of KCs and ALPNs is denoted by *M* and *N*, respectively. In this model, the PN activity is assumed to have zero mean, *r* = 0, and be uncorrelated, 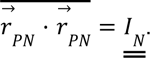

Here, *I* is an identity matrix and *r* denotes the average, *r*, is taken over independent realisations of *r_PN_*. Then, the element of the covariance matrix of *h* is

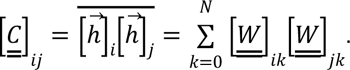

More detailed calculations can be found in (Litwin-Kumar et al. 2017). Randomised and homogeneous weights were used to populate *W*, such that each row in *W* has *K* elements that are 1 – α and *N* – *K* elements that are − α. The parameter α represents a homogeneous inhibition corresponding to the biological, global inhibition by APL. The value inhibition was set to be α = *A*/*M*, where *A* = 100 is an arbitrary constant and *M* is the number of KCs in each of the three datasets. The primary quantity of interest is the dimension of the KC activities defined by (Litwin-Kumar et al. 2017)

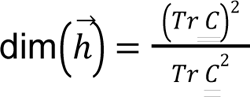

and how it changes with respect to *K*, the number of input connections. In other words, what are the numbers of input connections *K* onto individual KCs that maximise the dimensionality of their responses, ℎ, given *N* KCs, *M* ALPNs, and, a global inhibition α?

From Figure 5K, the theoretical values of *K* that maximise dim_(_ℎ_)_ in this simple model demonstrate the consistent shift towards lower values of FlyWire right datasets when compared with the hemibrain.

The limitations of the model are as follows:

*K* found in the FlyWire left and

1. The values in the connectivity matrix *W* take only two discrete values, either 0 and 1 or 1 – α and α. In a way this helps when calculating analytical results for the dimensionality of the KC activities. However, it is unrealistic as the connectomics data give the number of synaptic connections between the ALPNs and the KCs.
2. The global inhibition provided by APL to all of the mixing layer neurons is assumed to take a single value for all neurons. In reality, the level of inhibition would be different depending on the number of synapses between APL and the mixing layer neurons.
3. It is unclear whether the simple linear rate model presented in the original paper represents the behaviour of the biological neural circuit well. Furthermore, it remains unproven that the ALPN-KC neural circuit is attempting to maximise the dimensionality of the KC activities, albeit the theory is biologically well motivated (but _see_ 67,68_)._
4. The number of input connections to each mixing layer neuron is kept at a constant K for all neurons. It is definitely a simplification that can be corrected by introducing a distribution P(K) but this requires further detailed modelling.

### Mapping to the VirtualFlyBrain database

The VirtualFlyBrain (VFB) database^2^ curates and extracts information from all publications relating to *Drosophila* neurobiology, especially neuroanatomy. The majority of published neuron reconstructions, including those from the hemibrain, can be examined in VFB. Each individual neuron (i.e. one neuron from one brain) has a persistent ID (of the form *VFB_xxxxxxxx)*. Where cell types have been defined, they have an ontology ID (e.g. *FBbt_00047573*, the identified for the DNa02 descending neuron cell type). Importantly, VFB cross-references neuronal cell types across publications even if different terms were used. It also identifies driver lines to label many neurons. In this paper, we generate an initial mapping providing FBbt IDs for the closest and fine-grained ontology term that already exists in their database. For example, a FlyWire neuron with a confirmed hemibrain cell type will receive a FBbt ID that maps to that exact cell type while a descending neuron that has been given a new cell type might only map to the coarser term “adult descending neuron”. Work is already underway with VFB to assign both ontology IDs (FBbt) to all FlyWire cell types as well as persistent VFB_ ids to all individual FlyWire neurons.

## Acknowledgements and Contributions

We thank Andrew Champion and the MRC LMB Scientific Computing group for assistance with compute and web infrastructure. We thank Alex McLachlan, Rob Court, Claire Pilgrim, Damien Goutte-Gattat and David Osumi-Sutherland from the Virtual Fly Brain for helping mapping annotations into their ontology. We thank Forrest Collman and Casey Schneider-Mizell for developing and maintaining the CAVE engine and associated tools. We thank Ben Pedigo for discussions as well as help with matching and typing of some FlyWire neurons. Besides our own observations, we thank members of Rachel Wilson’s lab (ASB with Quinn Vanderbeck, Anna Li, Isabel Haber and Peter Gibb who reconstructed the AB neurons), Pratyush Kandimalla and Stephane Noselli for pointing out the apparent left/right inversion of FAFB and we thank Pratyush Kandimalla and Stephane Noselli for sharing their observations that situs inversus is extremely rare in wild-type *Drosophila*. ASB thanks Rachel I. Wilson for her support and interest as he finished this project after having moved to the Wilson lab; DH thanks Albert Cardona for support and mentoring while in his group. We thank Liqun Luo and Jakob Macke for comments on a previous version of this manuscript.

This work was supported by an NIH BRAIN Initiative grant 1RF1MH120679-01 to DB with GSXEJ; a Neuronex2 award to GSXEJ and DB (NSF 2014862, MRC MC_EX_MR/T046279/1); Wellcome Trust Collaborative Awards (203261/Z/16/Z and 220343/Z/20/Z) and core support from the MRC (MC-U105188491) to GSXEJ; EMBO fellowship (ALTF 1258-2020) and a Sir Henry Wellcome Postdoctoral Fellowship (222782/Z/21/Z) to ASB. For a full list of FlyWire Consortium members please see Dorkenwald *et al.^1^*.

